# Near-complete genome assembly of the African turquoise killifish enables locus-level profiling of transposable element expression

**DOI:** 10.64898/2026.10.12.757905

**Authors:** Yohey Ogawa, Akane Kawaguchi, Kota Abe, Tohru Ishitani, Shigehiro Kuraku

## Abstract

Tandem repeats (TRs) and transposable elements (TEs) occupy substantial fractions of eukaryotic genomes, yet how repeat detection programs affect downstream expression analyses remains poorly explored. Here, we present an improved chromosome-scale assembly of the African turquoise killifish, *Nothobranchius furzeri*. The assembly matches experimentally estimated genome size and chromosome number. It resolves highly expanded TRs that occupy up to 28.0% of the whole genome. Using this assembly and other complete vertebrate genome assemblies, we compared TR and TE-derived regions across combinations of TR detection programs and TE consensus library construction workflows under a standardized TE classification scheme. Choices in TR detection reshaped the apparent TE landscape. TE workflows shared a broad core of TE-derived regions but recovered complementary sets of age- and sex-associated TE loci, evaluated with a sex-balanced killifish multi-tissue aging cohort. Together, the combinatorial use of adaptable programs supports robust, locus-aware detection of complex repeats across genome assemblies.

## Introduction

The African turquoise killifish, *Nothobranchius furzeri*, has emerged as a non-mammalian vertebrate model for studying aging and adaptation to ephemeral habitats.^1–3^ Its naturally short lifespan and capacity for embryonic diapause allow researchers to investigate the genetic underpinnings of longevity and developmental arrest within a tractable timeframe. Furthermore, the phenotypic divergence between strains, such as the shorter-lived GRZ and the longer-lived MZM strains, has enabled the mapping of loci associated with lifespan.^4,5^ Unlocking the full potential of the killifish system requires a high-quality genomic reference.

The genomic architecture of the killifish poses substantial challenges for standard assembly strategies. Cytogenetic analyses have revealed a high abundance of repetitive sequences in the *N. furzeri* genome.^6^ A complete chromosome-scale assembly is therefore critical not only for accurately defining gene content but for resolving repeat landscapes associated with genomic evolution and structural variation. Previous efforts for sequencing the *N. furzeri* genome resulted in the acceptable reference genomes, MPIA-NFZ-2.0^7^, Nfu_20140520^8^, and NotFur1^5^, whose total assembly sizes are all no larger than 1.3 Gb and therefore below 1.5 Gb, an estimate from a cytogenetic report.^6^ More recently, Johnson et al.^9^ reported UI_Nfuz_MZM_1.0, a 1.5-Gb assembly. However, this assembly consists of more than 2,600 non-chromosomal scaffolds and includes 24 chromosome-scale sequences, exceeding the chromosome number estimated from cytogenetic reports (n = 19).^6,10^ UI_Nfuz_MZM_1.0 also recovers no more than 97% of the reference genes in conventional gene completeness assessment (Figure 1A). A more recent assembly did not reconstruct chromosomal sequences, but the total assembly size resembles the estimated genome size.^11^ The discrepancy in chromosome numbers and the modest gene space coverage preclude precise genome-wide analyses. These limitations require further attempts to establish a definitive, high-quality reference genome.

**Figure 1.**
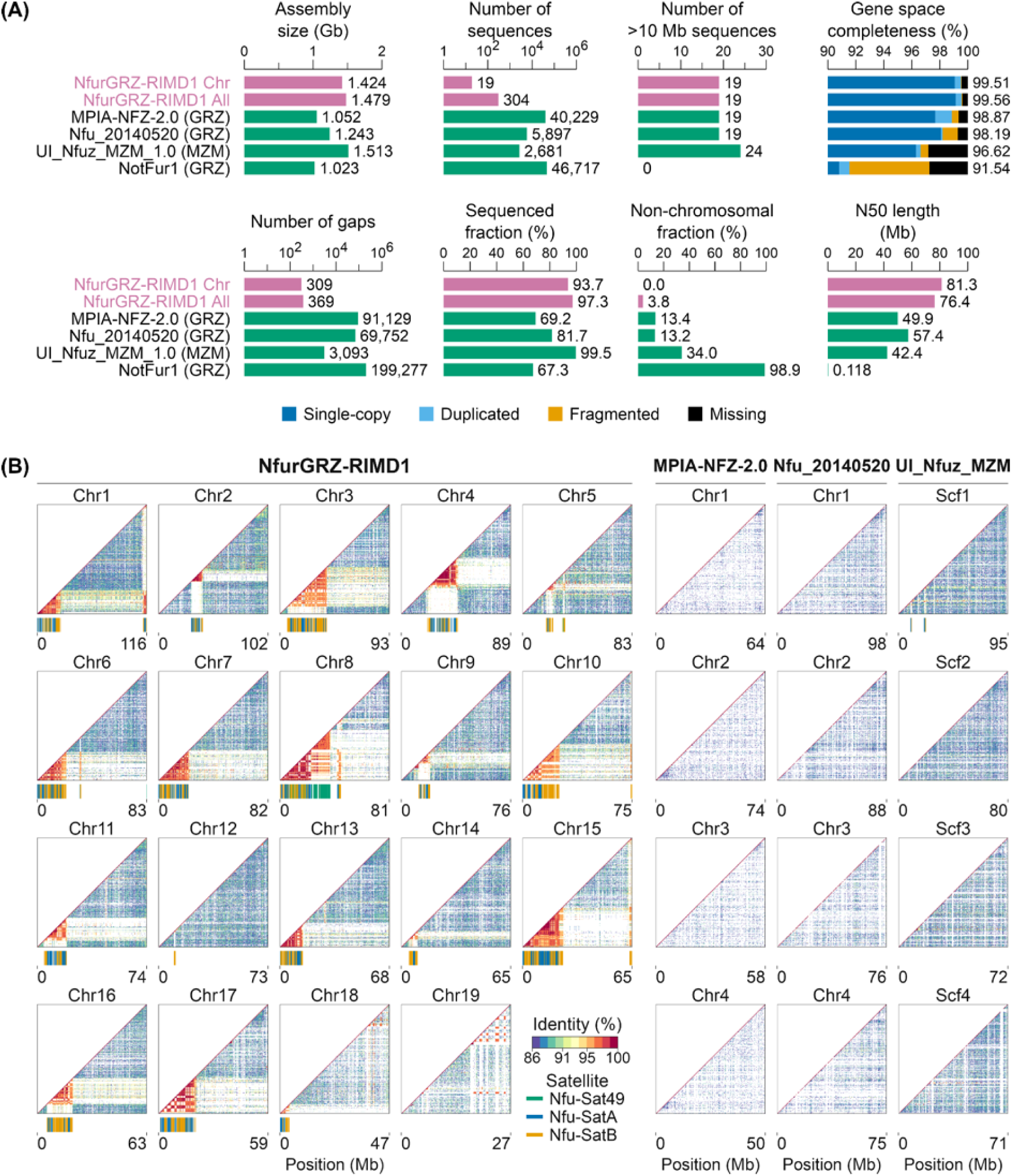
Chromosome-scale assembly and satellite-rich regions of the killifish genome. (A) Assembly metrics for the 19 chromosome-scale sequences of NfurGRZ-RIMD1 (“Chr”) and the full assembly including unplaced scaffolds (“All”), shown in magenta. Metrics for four previously published N. furzeri assemblies are shown in green for comparison: MPIA-NFZ-2.0^7^; Nfu_20140520^8^; UI_Nfuz_MZM_1.0^9^; and NotFur1^5^. (B) Lower-triangular self-similarity dot plots for the 19 chromosome-scale sequences of NfurGRZ-RIMD1, four chromosome-scale sequences each from MPIA-NFZ-2.0 and Nfu_20140520, and four scaffolds from UI_Nfuz_MZM_1.0. Colors indicate pairwise sequence identity, and sequence lengths are shown in megabases below each plot. Tracks beneath each plot indicate matches to the previously reported centromeric satellite repeat units Nfu-Sat49, Nfu-SatA, and Nfu-SatB. Only matches meeting the specified criteria are shown (Methods). Chr, chromosome; Scf, scaffold.

Transposable elements (TEs) are widespread components of eukaryotic genomes and contribute to genome evolution.^12,13^ A comprehensive genome-wide repeat landscape requires a high-quality collection of consensus TE sequences. A TE library, containing such consensus sequences in a selected genome, should ideally be manually constructed. TE library construction is thus labor-intensive and requires specialized expertise, making it infeasible to readily keep pace with the rapid accumulation of new genome assemblies. In addition, a comprehensive view of genomic repeats must include tandem repeats (TRs), in which identical or similar DNA sequence units are repeated sequentially. TRs include simple repeats composed of short sequence motifs and satellite DNA that can form extensive arrays, such as those found at centromeres.^14^ The wide variation in repeat-unit length and total array size challenges comprehensive TR detection across the genome.^15^ Automated workflows continue to be developed for TR detection, *de novo* TE consensus construction, TE library curation, and genome-wide identification of TE-derived regions.^15–20^ Previous studies have repeatedly benchmarked TE discovery workflows^21–23^ and compared TR detection programs.^24^ Yet, how different combinations of TR detection programs and automated TE consensus library workflows affect the identification of TE-derived genomic regions remains poorly explored.

How can we select an appropriate workflow for identifying TE-derived regions across the genome of a given species? TE content and composition vary both among and within vertebrate lineages.^25^ Such variation potentially limits the applicability of benchmark results across species.^26,27^ In addition, benchmarks have relied on TE regions derived from manually curated libraries in a limited set of species.^21^ A complementary approach is to assess how workflow choice affects downstream analyses that use the resulting TE-derived genomic regions. Locus-resolved analysis of age- and sex-associated TE expression provides one such application. LINE-1 expression patterns vary with age and sex^28^, and TE derepression has been linked to cellular dysfunction during aging.^29–31^ In the African turquoise killifish, previous work has quantified age- and sex-associated TE expression by aggregating transcript counts across genomic copies assigned to the same TE consensus sequence.^32^ Analysis at individual TE loci can localize TE expression signals to specific genomic regions, but locus-resolved analyses depend on the TE genomic intervals identified by each detection workflow. Comparing TE loci with age- or sex-associated expression signals across workflows can thus reveal both shared findings and workflow-dependent differences.

Here, we present a substantially improved chromosome-scale assembly of the African turquoise killifish GRZ strain. The assembly closely matches the experimentally estimated genome size, contains the expected number of chromosome-scale sequences, and has substantially fewer gaps than previous reference assemblies. The assembly also better represents TRs, including centromeric satellite-rich regions. We compared genomic coverage and interval-length distributions of TRs identified by four detection programs across five vertebrate genomes. Focusing on three teleost fish genomes, we generated TE consensus libraries from eight automated workflows. After classifying TEs with a uniform classification method, we detected TR and TE-derived regions across all 32 TR-TE program combinations. Finally, we quantified expression signals at individual TE loci in a sex-balanced, multi-tissue aging cohort of the killifish and compared TE loci showing age- and sex-associated expression signals across TE workflows.

## Results

### Highly repetitive *N. furzeri* genome assembly

In the present study, we sequenced the genome of the African turquoise killifish GRZ strain, which we hereafter refer to as killifish. Muscle tissue from an adult male was used for high-fidelity (HiFi) long-read sequencing and assembly followed by Hi-C scaffolding (Methods). This process yielded a chromosome-level genome assembly, NfurGRZ-RIMD1, with the total size of 1.48 Gb and 19 chromosome-scale sequences (Figure 1A). The assembly size is close to the previously estimated genome size of 1.52 Gb, and the number of chromosome-scale sequences matches the reported haploid chromosome number.^6^ The numbers and fractions of undetermined regions (‘gaps’) as well as non-chromosomal sequences have been dramatically reduced (Figure 1A). More than 99.5% of the single-copy ortholog markers were classified as “complete” in this assembly (Methods), indicating high completeness of gene content.

To evaluate the representation of repetitive pericentromeric and centromeric regions in the newly assembled killifish genome, we identified tandem repeat intervals and their consensus repeat unit sequences utilizing the TRASH program.^33^ We therein projected three centromeric repeat units, Nfu-Sat49, Nfu-SatA, and Nfu-SatB, previously documented.^6,34^ We identified satellite-rich regions on all assembled chromosomes except chromosome 19, and these regions contained repeats highly similar to previously reported centromeric satellites (Figure 1B). In contrast, two previous genome assemblies (MPIA-NFZ-2.0 and Nfu_20140520) of this species lack clearly identifiable centromeric repeat regions, whereas UI_Nfuz_MZM_1.0 contains centromeric repeats mainly on short scaffolds (Figure S1). These results suggest that our assembly, NfurGRZ-RIMD1, achieves genome-wide recovery of repeat-rich regions previously observed cytogenetically.

### Parameter-dependent variation in repeat regions

To examine whether tandem repeats are abundant in the killifish genome, we compared their abundance by repeat unit length across nearly complete vertebrate genome assemblies (zebrafish^35^, medaka^36^, mouse^37^, and human^38,39^). After clustering the repeat unit sequences identified by TRASH, we found that tandem repeats with units of 77 bp and 348 bp each account for more than 7% of the killifish genome (Figure 2A). Similarly, in zebrafish, tandem repeats with repeat unit lengths of 92 bp and 186 bp each account for more than 2% of the genome. In contrast, medaka, mouse, and human showed lower tandem repeat abundances and relatively broad distributions of repeat unit lengths. The most abundant tandem repeats in killifish include those with repeat units longer than 300 bp, whereas those in the other species generally have repeat units shorter than 200 bp. These results suggest a pronounced expansion of tandem repeats with long repeat units in the killifish.

**Figure 2.**
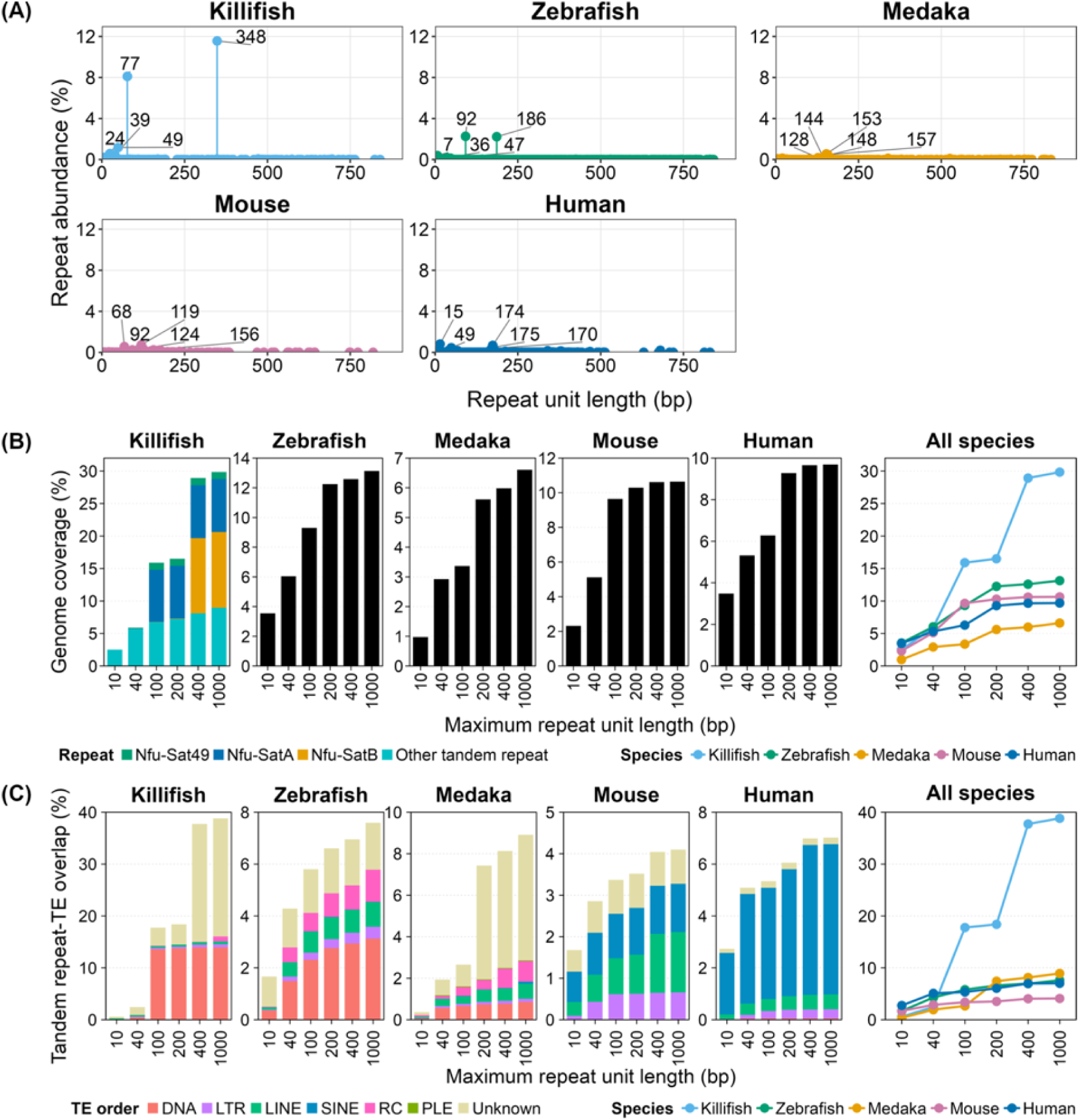
Tandem repeat profiles and parameter-dependent overlap with TE-derived regions. (A) TRASH-derived repeat unit landscapes in killifish, zebrafish, medaka, mouse, and human. Similar repeat unit sequences were clustered as described in Methods, and the abundances of clusters with the same representative repeat unit length were summed. Each point shows the representative repeat unit length in base pairs (x-axis) and the corresponding total genomic abundance of tandem repeats, expressed as a percentage of genome size (y-axis). (B) Tandem repeat coverage obtained with TRF at maximum repeat unit length settings of 10, 40, 100, 200, 400, or 1,000 bp, expressed as a percentage of each genome. Stacked segments distinguish TRF intervals overlapping genomic matches to the centromeric satellite repeat units Nfu-Sat49, Nfu-SatA, or Nfu-SatB from all other tandem repeat intervals. (C) Overlap between TE-derived genomic regions and the TRF intervals in (B). TE-derived regions were identified with RepeatMasker using RepeatModeler2 libraries. The total height of each bar shows the percentage of all TE-derived bases overlapped by TRF intervals, and stacked segments show the contribution of each TE order to the overlap. RC, rolling-circle transposable element; PLE, Penelope-like transposable element; Unknown, no TE order assigned.

The high abundance of tandem repeats with long repeat unit lengths in killifish prompted us to evaluate how program choice affects repeat profiling. Tandem Repeats Finder (TRF^40^) is incorporated into standard workflows for detecting TE-derived genomic regions (e.g., RepeatMasker) and is one of the most widely used programs for automatically detecting tandem repeats. We systematically varied the maximum repeat unit length parameter in TRF and quantified the tandem repeat fractions across the five species’ genomes (Figure 2B). We first set a maximum repeat unit length of 10 bp because it is the value used in RepeatMasker’s initial stage of tandem repeat detection. In killifish, tandem repeat coverage was less than 5% of the assembly with the initial TRF setting. Tandem repeat coverage reached approximately 30% as the maximum repeat unit length parameter increased. Marked increases in coverage occurred at settings of 100 bp and 400 bp. The step increases were consistent with the high abundance of repeats with unit lengths of 77 bp and 348 bp identified by TRASH (Figure 2A). In addition, a larger fraction of TE-derived genomic regions was overlapped by tandem repeats as the maximum repeat unit length increased (Figure 2C). The overlap was particularly pronounced for DNA transposon and LTR retrotransposon categories (Figure S2). The other species also showed smaller increases in tandem repeat coverage and overlap with TE-derived regions, with the increase in overlap particularly evident for LINE retrotransposons in zebrafish and SINE retrotransposons in mouse and human (Figures 2B, 2C, and S2). Together, these results imply that insufficient masking of tandem repeats with long repeat units can inflate estimates of TE genomic coverage. The choice of tandem repeat detection parameters substantially affects TE-derived region detection, especially in genome assemblies rich in tandem repeats with long repeat units.

### Broad coverage of tandem repeat regions by ULTRA

To assess how program choice affects tandem repeat detection, we compared four programs: the widely used programs, TRF and tantan^41^, as well as the more recently developed programs, ULTRA^15^ and longdust^24^. We used near-complete genome assemblies of five species with distinct tandem repeat profiles: killifish, zebrafish, medaka, mouse, and human. We examined whether a single program could comprehensively detect both short-period tandem repeat intervals and long-period satellite-like tandem repeat arrays. To compare the four detection programs across the five species, we plotted cumulative tandem repeat coverage against interval length (Figure 3A). ULTRA yielded the greatest tandem-repeat coverage among the programs in all species analyzed. The largest fraction of tandem repeat coverage consisted of bases detected by all four programs, followed by bases detected only by ULTRA (Figure 3B). The other three programs produced comparable total tandem repeat coverage, although their repeat interval length distributions differed substantially (Figure 3A). Compared with TRF and longdust, tantan showed a larger contribution of short intervals to its total tandem repeat coverage. Consistent with these differences, pairwise comparisons showed that the fraction of detected tandem repeat bases covered by the other two programs was generally lower for tantan than for TRF or longdust (Figure 3C). In contrast, ULTRA detected tandem repeat intervals across the broadest range of lengths (Figure 3A). These intervals covered a large fraction of tandem repeat bases detected by each of the other three programs (Figure 3C). These trends were consistent across all five genomes. Based on ULTRA’s broad tandem repeat coverage, we selected it to more comprehensively detect repeat intervals for downstream analyses.

**Figure 3.**
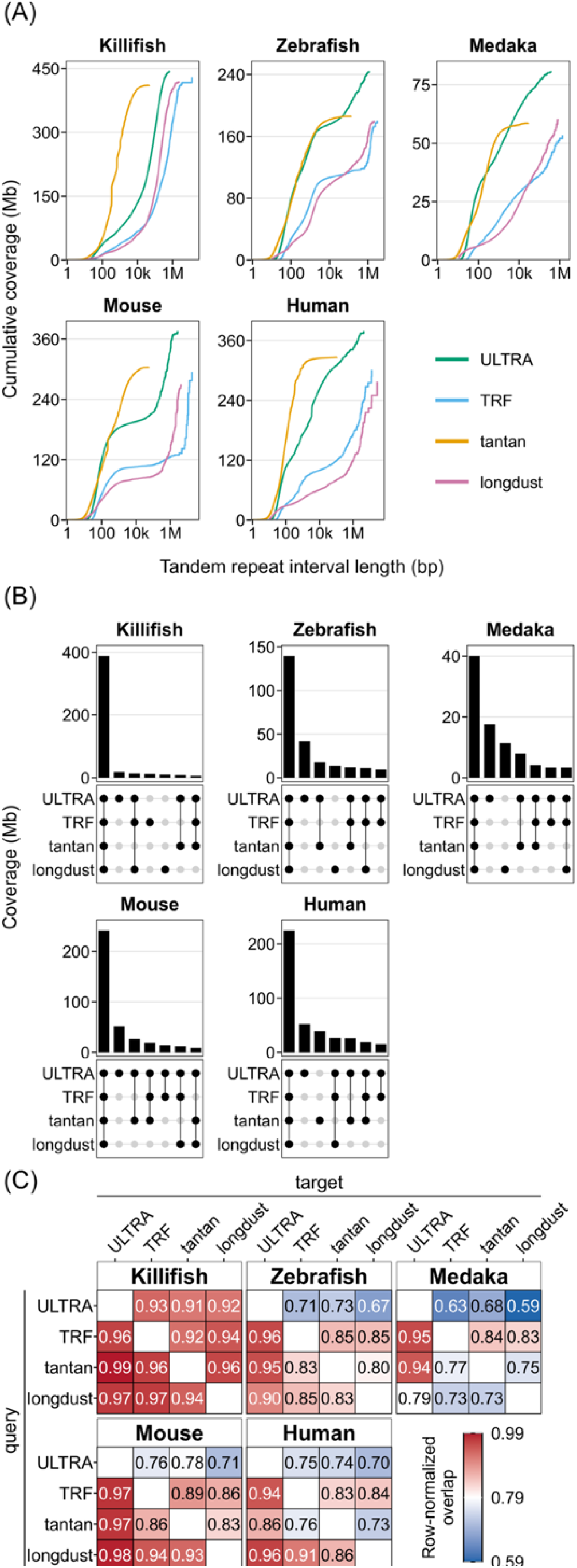
Comparison of tandem repeat coverage across four detection programs. (A) Cumulative genomic coverage of merged tandem repeat intervals detected by tantan, TRF, ULTRA, and longdust in killifish, zebrafish, medaka, mouse, and human. For each species and program, merged intervals were ordered from shortest to longest, and their lengths were cumulatively summed. The x-axis shows merged interval length on a log10 scale, and the y-axis shows cumulative coverage in megabases. (B) UpSet plots showing the seven largest mutually exclusive base-pair intersections among the four programs for each species. Each bar shows the total genomic coverage of bases detected by exactly the indicated combination of programs. (C) Pairwise overlap of tandem repeat regions at base-pair resolution. For each species, the cell in row X and column Y shows the fraction of tandem repeat bases detected by program X that are also detected by program Y. Values are shown in the cells and represented by color.

### TE order classification through majority voting

To characterize genomic repeat landscapes, we designed a custom pipeline for tandem repeat detection, TE consensus library construction, TE classification, and genome-wide detection of TE-derived regions (Figure 4A). We compared state-of-the-art programs for key pipeline steps using vertebrate genome assemblies. First, we evaluated TE classification programs using curated TE consensus sequences and their reference classification labels from Dfam, a public database of TE families.^42^ In the present study, we used six harmonized TE categories, hereafter referred to as TE orders: DNA, LTR, LINE, SINE, RC, and PLE. We classified the curated TE consensus sequences from zebrafish, mouse, and human using four machine learning classifiers: Terrier^43^, CREATE^44^, NeuralTE^45^, and TEclass2^46^. All four classifiers showed high agreement with the Dfam reference order labels, except for TEclass2 in zebrafish (Figure 4B). To extend classifier comparison to *de novo* TE consensus sequences, we selected killifish, zebrafish, and medaka. These species represent lineages separated by more than 200 million years of evolution^47^, and their TE compositions are therefore expected to differ substantially. For each species, four TE consensus libraries were generated using Earl Grey^19^, RepeatModeler2^16^, EDTA^21^, and HiTE^20^. Four additional libraries were obtained by refining the RepeatModeler2- and EDTA-built libraries separately with the TE library curation programs, MCHelper^17^ or TEtrimmer^18^, producing eight libraries per species (Methods). The resulting TE consensus sequences were classified using the four machine learning classifiers and RepeatClassifier, a current standard homology-based classification method that uses BLAST searches.^16^ We then calculated pairwise exact-match rates for the assigned TE order labels. Median pairwise exact-match rates across the 24 combinations (3 species and 8 libraries) were generally higher between RepeatClassifier and the machine learning classifiers (0.55-0.59) than among the machine learning classifiers (0.47-0.59; Figure 4C). NeuralTE and TEclass2 generally showed relatively low agreement with the other machine learning classifiers (Figure S3). Together, these comparisons showed that TE order assignments to *de novo* consensus sequences depended substantially on classifier choice.

**Figure 4.**
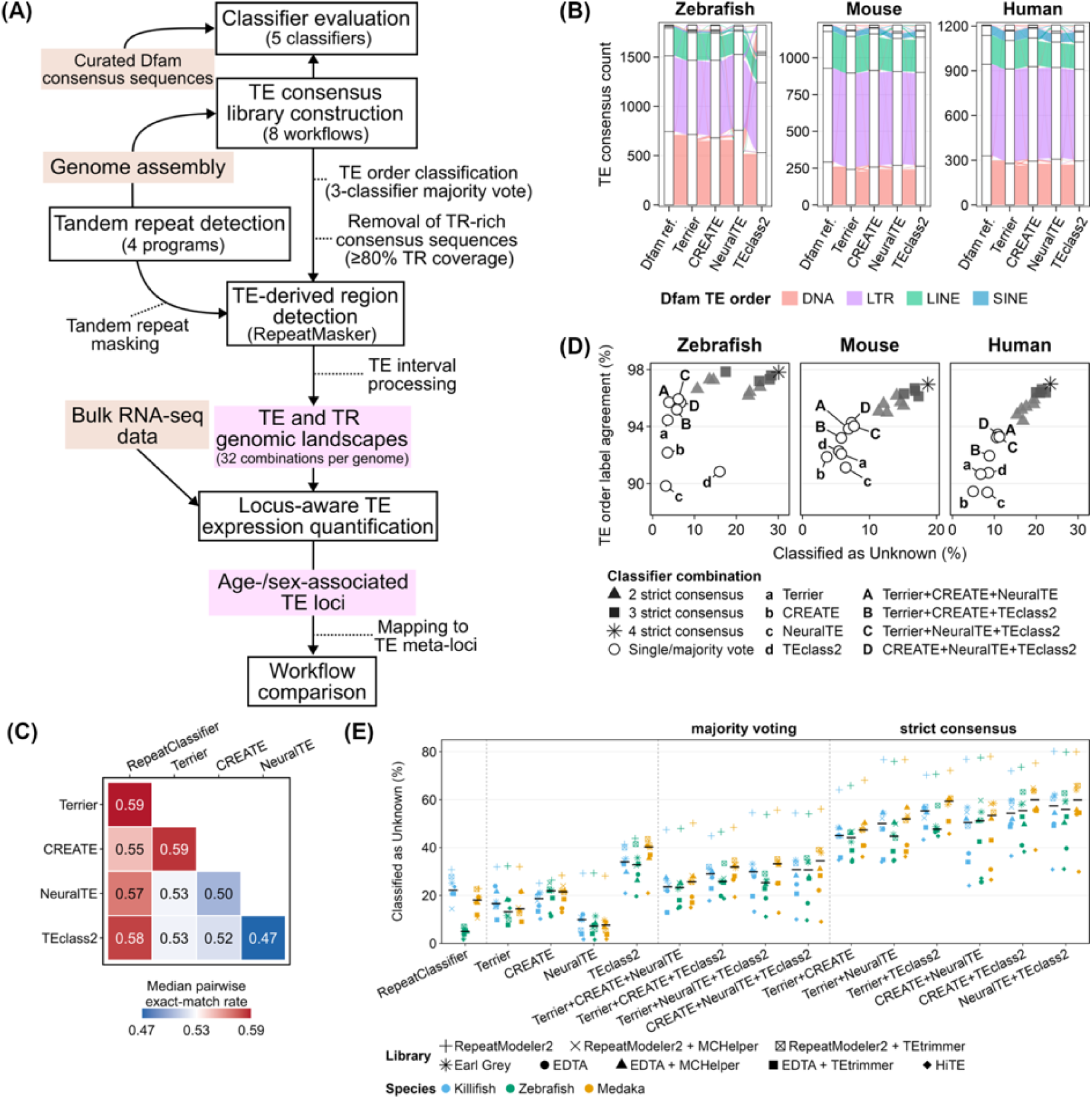
Analysis workflow and evaluation of TE order classification. (A) Overview of the analysis workflow. (B) Alluvial plots comparing Dfam reference TE order assignments with those made by Terrier, CREATE, NeuralTE, and TEclass2 for curated TE consensus sequences from zebrafish, mouse, and human. Ribbon colors indicate the Dfam reference TE order. (C) Pairwise exact-match rates of TE order assignments among RepeatClassifier and the four machine learning classifiers for *de novo* TE consensus sequences. Each cell shows the median across 24 combinations of three species (killifish, zebrafish, and medaka) and eight libraries. See also Figure S3. (D) Comparison of individual classifiers and classifier combinations using curated Dfam TE consensus sequences with known TE order labels. Each point represents an individual classifier or a combination of classifiers using strict consensus or majority voting, as indicated in the key. The x-axis shows the percentage of all evaluated sequences labeled “Unknown”. The y-axis shows agreement with Dfam, calculated as the percentage of sequences assigned the same TE order as the reference after excluding sequences labeled “Unknown”. (E) “Unknown” assignment rates for eight *de novo* TE consensus libraries from each of three species: killifish, zebrafish, and medaka. Sequences were classified using individual classifiers or by combining classifier predictions using strict consensus or majority voting. Each point represents one species-library combination. Point shape indicates the library construction workflow, and color indicates the species. For each classification strategy on the x-axis, black horizontal bars indicate the median across the eight libraries within each species.

Machine learning classifiers can complement alignment-based classification by recognizing patterns learned from TE sequences.^43,44^ To test whether combining predictions from multiple machine learning classifiers could improve TE order classification, we compared two strategies: “strict consensus” and “majority vote” (Methods). Sequences were labeled “Unknown” when no TE order met the agreement criterion for a given strategy. Using Dfam reference TE consensus sequences, we compared label agreement and “Unknown” assignment rates across classification strategies (Figure 4D). Both strict-consensus and majority-vote strategies improved agreement with the reference order labels in zebrafish, mouse, and human. However, strict consensus substantially increased the fraction of sequences labeled “Unknown”, whereas three-classifier majority voting resulted in only a modest increase (Figure 4D). These results support three-classifier majority voting as a practical approach for improving agreement with the reference labels while limiting the increase in “Unknown” assignments.

We next evaluated these strategies on the *de novo* TE consensus libraries generated from the killifish, zebrafish, and medaka genome assemblies (Figure 4E). In zebrafish, RepeatClassifier yielded a median “Unknown” rate of 5.0% across the eight libraries. In contrast, species-specific curated Dfam libraries were unavailable for killifish and medaka at the time of this study, and median “Unknown” rates from RepeatClassifier were 22.2% and 18.1%, respectively, across the eight libraries generated for each species. For the four three-classifier majority-vote combinations, the corresponding median rates were as follows (killifish and medaka, respectively): 23.6% and 25.7% (Terrier + CREATE + NeuralTE); 29.0% and 31.9% (Terrier + CREATE + TEclass2); 29.9% and 33.2% (Terrier + NeuralTE + TEclass2); and 30.8% and 34.4% (CREATE + NeuralTE + TEclass2). In addition, the proportions of TE orders assigned by majority voting were broadly similar to those obtained with RepeatClassifier (Figure S4). Together, these results suggest that three-classifier majority voting is a practical approach to classify *de novo* TE consensus sequences at the order level. Because TEclass2 generated higher “Unknown” rates than the other individual classifiers (Figure 4E) and showed relatively low agreement with the other machine learning-based classifiers (Figure 4C), we selected Terrier, CREATE, and NeuralTE for majority voting in downstream analyses.

### Variation in TE consensus sequence number and length across workflows

We generated *de novo* TE consensus libraries from the killifish, zebrafish, and medaka genomes using eight workflows described in the preceding section. The resulting consensus sequences were reclassified at the TE-order level using our majority-vote classification strategy. We then compared the number, sequence-length summaries, and GC-content of consensus sequences across workflows (Figure 5). The number of consensus sequences varied substantially among workflows (Figure 5A). In zebrafish, for which a Dfam reference library was available for comparison, the *de novo* libraries generally contained more DNA transposon consensus sequences but fewer LTR, LINE, and SINE retrotransposon consensus sequences than the reference library. Among the *de novo* pipelines, Earl Grey generated the largest numbers of LINE and PLE consensus sequences across all three species. EDTA generated more than twice as many DNA transposon consensus sequences as the other pipelines in all three species, as well as a relatively large number of LTR consensus sequences.

**Figure 5.**
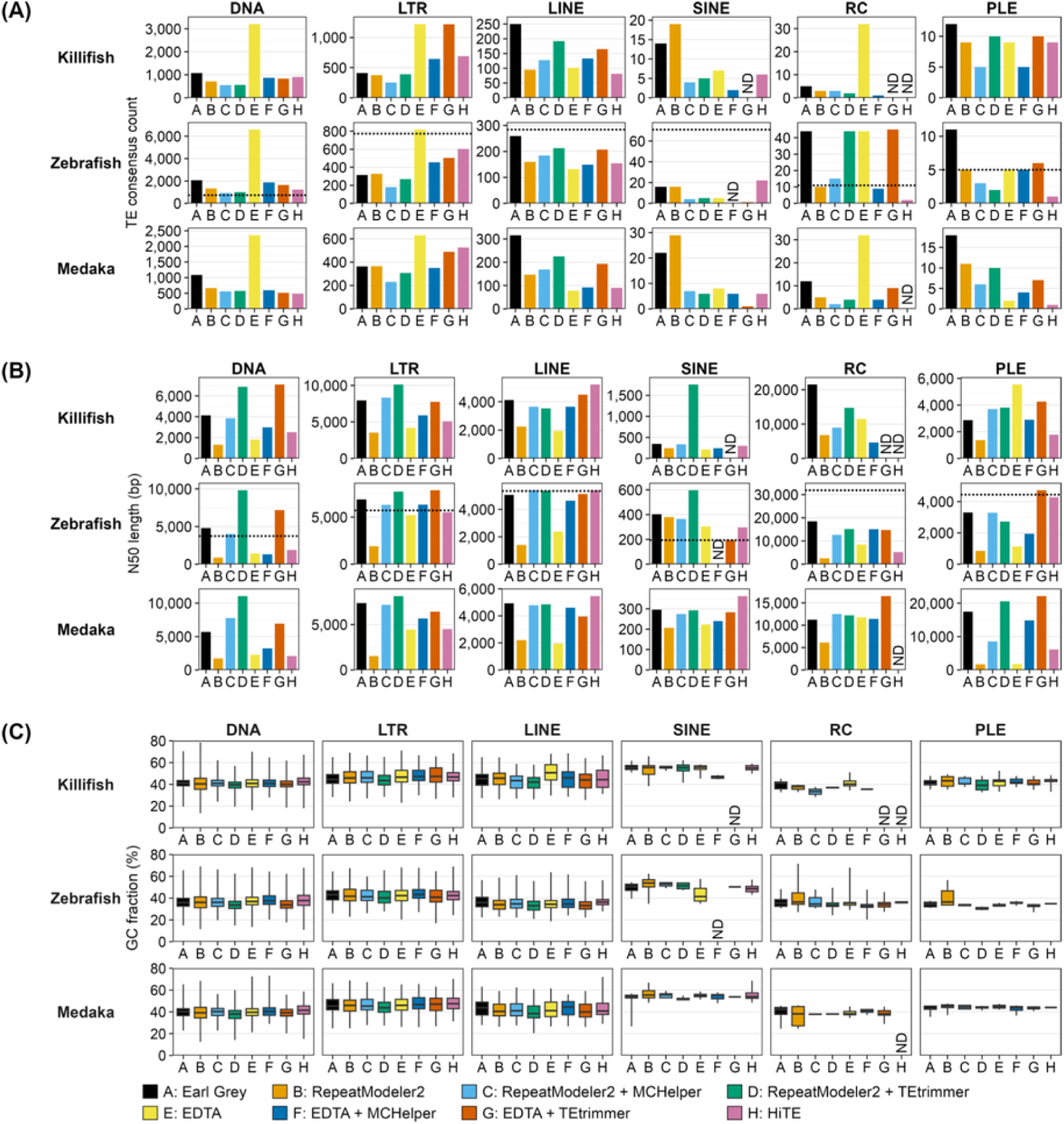
Characteristics of TE consensus sequences across library construction workflows. (A) Number of TE consensus sequences and (B) N50 sequence length for killifish, zebrafish, and medaka, stratified by TE order and library construction workflow. Rows represent species, and columns represent TE orders. In the zebrafish row, black dotted lines indicate the corresponding values for the Dfam reference library. (C) Box plots showing the GC-content distributions of TE consensus sequences. Boxes span the first to third quartiles, center lines indicate medians, and whiskers show the minimum and maximum values. Black dotted lines in the zebrafish row indicate the median GC-content of Dfam consensus sequences within each TE order. Consensus sequences classified as “Unknown” were excluded from the analyses. ND indicates that no consensus sequence was assigned to the corresponding species–order–workflow combination.

Consensus sequence lengths differed markedly among workflows. In the zebrafish RepeatModeler2-built library, TE consensus sequences were generally shorter than those in the Dfam reference library except for SINE retrotransposons (Figure 5B). Similarly, the shorter consensus sequence lengths were observed in the EDTA-built library for DNA, LINE, RC, and PLE transposons. By contrast, consensus sequence lengths in the HiTE and Earl Grey libraries were broadly comparable to those in Dfam, except for RC transposons. Comparisons among workflows in killifish and medaka also showed that RepeatModeler2 and EDTA tended to yield shorter consensus sequences. Refinement with MCHelper extended sequence lengths broadly comparable to those in Dfam for LTR and LINE retrotransposons in zebrafish. TEtrimmer also increased the lengths of EDTA- and RepeatModeler2-derived sequences, although some resulting sequences were longer than Dfam consensus sequences within the same TE order. In addition, refinement with MCHelper or TEtrimmer reduced the number of DNA transposon consensus sequences and increased that of LINE consensus sequences in both the EDTA and RepeatModeler2 libraries across all three species (Figure 5A). In zebrafish, these changes brought the consensus sequence counts closer to those in the Dfam reference library. The GC-content of TE consensus sequences was broadly similar among libraries (Figure 5C). Overall, both the number and length of TE consensus sequences were workflow-dependent, with subsequent refinement generally yielding libraries with longer consensus sequences.

### Highest combined TE and TR coverage with Earl Grey-ULTRA

We compared the TE-derived genomic regions identified using the eight consensus libraries in each of the three fish genomes. To limit potential overestimation of TE coverage in regions overlapping TRs, we first masked TR regions using one of the four TR detection programs. We then identified TE-derived regions with RepeatMasker using each of the eight TE consensus libraries, yielding 32 combinations per species (Methods and Figure 4A). TE and TR coverage was calculated from processed repeat intervals after fragment joining, overlap filtering, and exon exclusion (Methods). Among the TR detection programs, ULTRA yielded the highest TR coverage in all three species after integration with TE-derived regions and subsequent repeat-track processing (median across the eight workflows: 27.7% in killifish, 13.2% in zebrafish, 6.8% in medaka; Figure 6A), consistent with our earlier comparison of TR detection programs (Figure 3). Following TR masking, TE coverage was highest with the Earl Grey-built library in all three species (median across the four TR programs: 40.3% in killifish, 48.3% in zebrafish, and 35.0% in medaka). Among the 32 combinations examined, Earl Grey-ULTRA yielded the highest combined TE and TR coverage in all three fish genomes (67.8% in killifish, 60.3% in zebrafish, and 41.4% in medaka).

**Figure 6.**
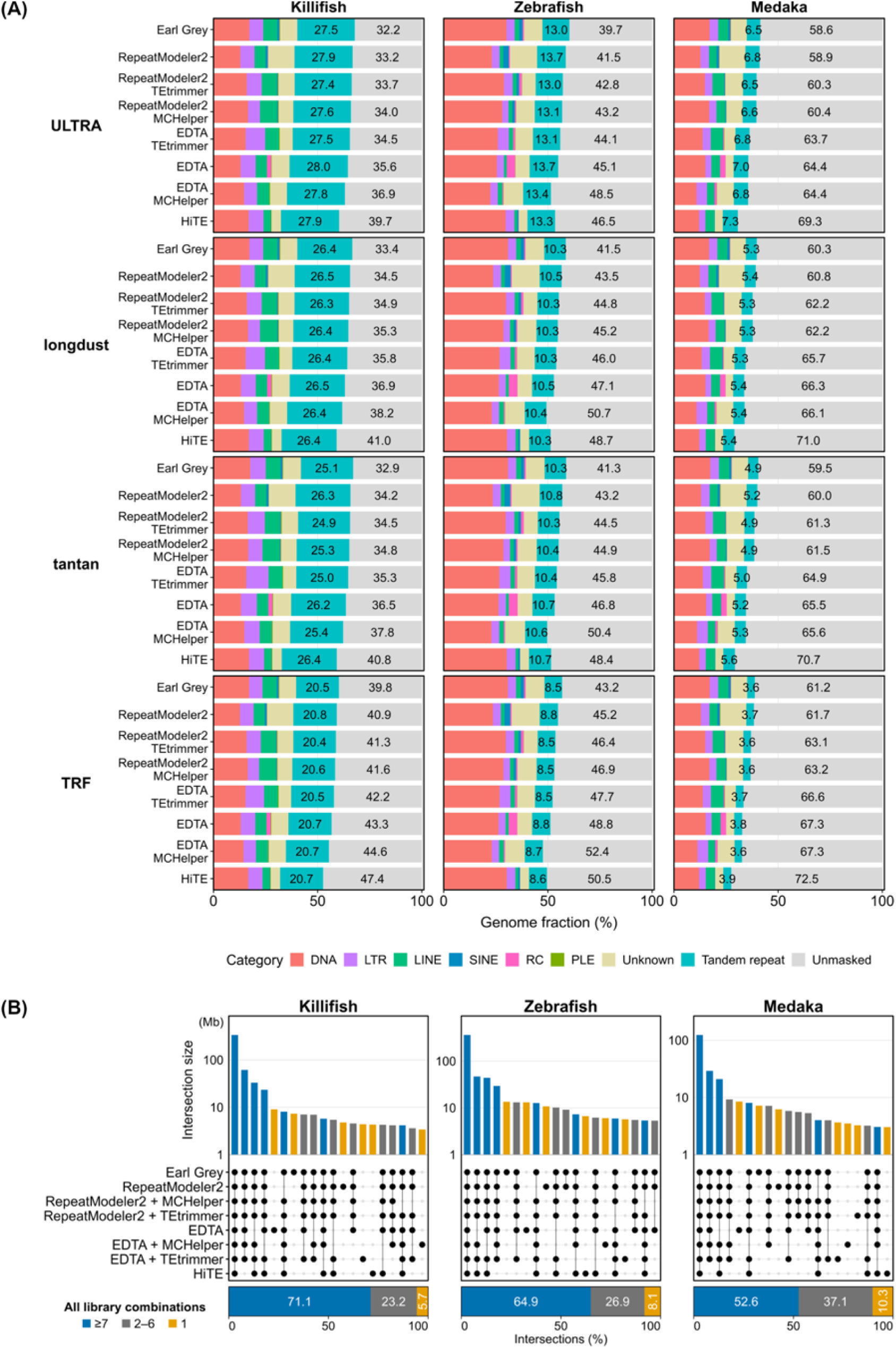
Genome-wide repeat coverage and overlap among library-defined TE regions. (A) Genome-wide repeat coverage in killifish, zebrafish, and medaka for all 32 combinations of four tandem repeat masking methods and eight TE library construction workflows. Libraries are shown in the same order across panels, arranged by increasing unmasked fraction in killifish with ULTRA masking. Stacked segments show the percentage of each genome assigned to each TE order, TEs classified as “Unknown”, tandem repeats, or unmasked sequence. The unmasked coverage was the portion of the genome not covered by any of the repeat intervals. (B) Upper UpSet plots show the 20 largest mutually exclusive base-pair intersections among the eight TE tracks for each species, using the ULTRA tandem repeat mask. Each bar shows the total length, in megabases, of genomic regions identified as TE-derived using exactly the libraries indicated by the connected-dot matrix. The y-axis is logarithmic. Lower stacked bars summarize all library combinations, including those not shown in the UpSet plots, as percentages of the union of TE-derived regions across the eight libraries. In both plots, blue denotes regions identified using seven or eight libraries, orange denotes regions identified using a single library, and gray denotes regions identified using two to six libraries.

We next compared the TE-derived regions among the eight libraries at base-pair resolution, with TR masking fixed to ULTRA (Figure 6B). For each species, we calculated the union of TE-derived regions across the eight libraries and counted the number of libraries that identified each genomic position as TE-derived. TE-derived genomic positions identified using seven or eight libraries accounted for 71.1% of the union in killifish, 64.9% in zebrafish, and 52.6% in medaka. Each library also identified genomic positions not detected as TE-derived using any of the other seven libraries. Among the eight libraries, EDTA and Earl Grey contributed relatively large unique fractions. TE-derived genomic positions identified only with EDTA accounted for 1.35%, 1.64%, and 2.29% of the union in killifish, zebrafish, and medaka, respectively. The corresponding fractions for positions identified only with Earl Grey were 1.09%, 1.70%, and 1.95%. Overall, the Earl Grey-ULTRA combination consistently yielded the greatest genome-wide repeat coverage together with appreciable library-dependent TE coverage.

### Variation in TE sequence divergence across workflows

We quantified nucleotide sequence divergence between genomic TE copies and their assigned consensus sequences using RepeatMasker alignments (Figure S5A). TE-derived regions identified with the EDTA-built library showed higher sequence divergence than those generated with the other libraries. Refinement of the EDTA consensus sequences with TEtrimmer or MCHelper shifted the distributions toward those observed for the other libraries. In contrast, the HiTE-built library yielded narrower distributions concentrated at low divergence values in all three species. This library generally yielded the lowest combined TE and TR coverage (Figure 6A). The RepeatModeler2-built library, its refined versions, and the Earl Grey-built library yielded divergence distributions intermediate between those obtained with EDTA and HiTE (Figure S5A). We observed similar divergence patterns when the analysis was restricted to TE-derived regions overlapping genomic intervals identified across all eight libraries (Figure S5B).

### Tissue-dependent chromosomal distributions of TE loci with age- and sex-associated expression levels

We examined how expression at TE loci varied among tissues and with age and sex. The TE track generated using the Earl Grey-built library after ULTRA tandem repeat masking was used for the following expression and chromosomal distribution analyses because the Earl Grey-ULTRA combination yielded the highest combined TE and TR coverage (Figure 6). Expression of genes and individual TE loci was quantified using the TE-Seq workflow^48^ with bulk transcriptome data from a multi-tissue killifish aging cohort.^49^ We retained nine tissues for the following analyses after excluding sex-specific organs and tissues with limited biological replication or low-quality RNA (Methods). To examine variation in expression patterns, we conducted principal component analysis (PCA) separately for gene expression and TE locus expression, both across all nine tissues and within each tissue (Figure 7A). Across the nine tissues analyzed, the gene-based PCA showed clear separation of samples by tissue (Figure 7A, upper-left panel). Similarly, PCA using only TE locus expression showed overall tissue-level separation, although samples from several tissues partially overlapped. Within individual tissues, both gene-based PCA and TE-based PCA showed clear separation between female and male samples in the kidney, liver, and caudal fin (Figure 7A). Notably, brain and spinal cord samples showed clearer sex-associated separation in the TE-based PCA than in the corresponding gene-based PCA. Age-associated ordering of samples was most evident in muscle in both the gene- and TE-based PCA projections.

**Figure 7.**
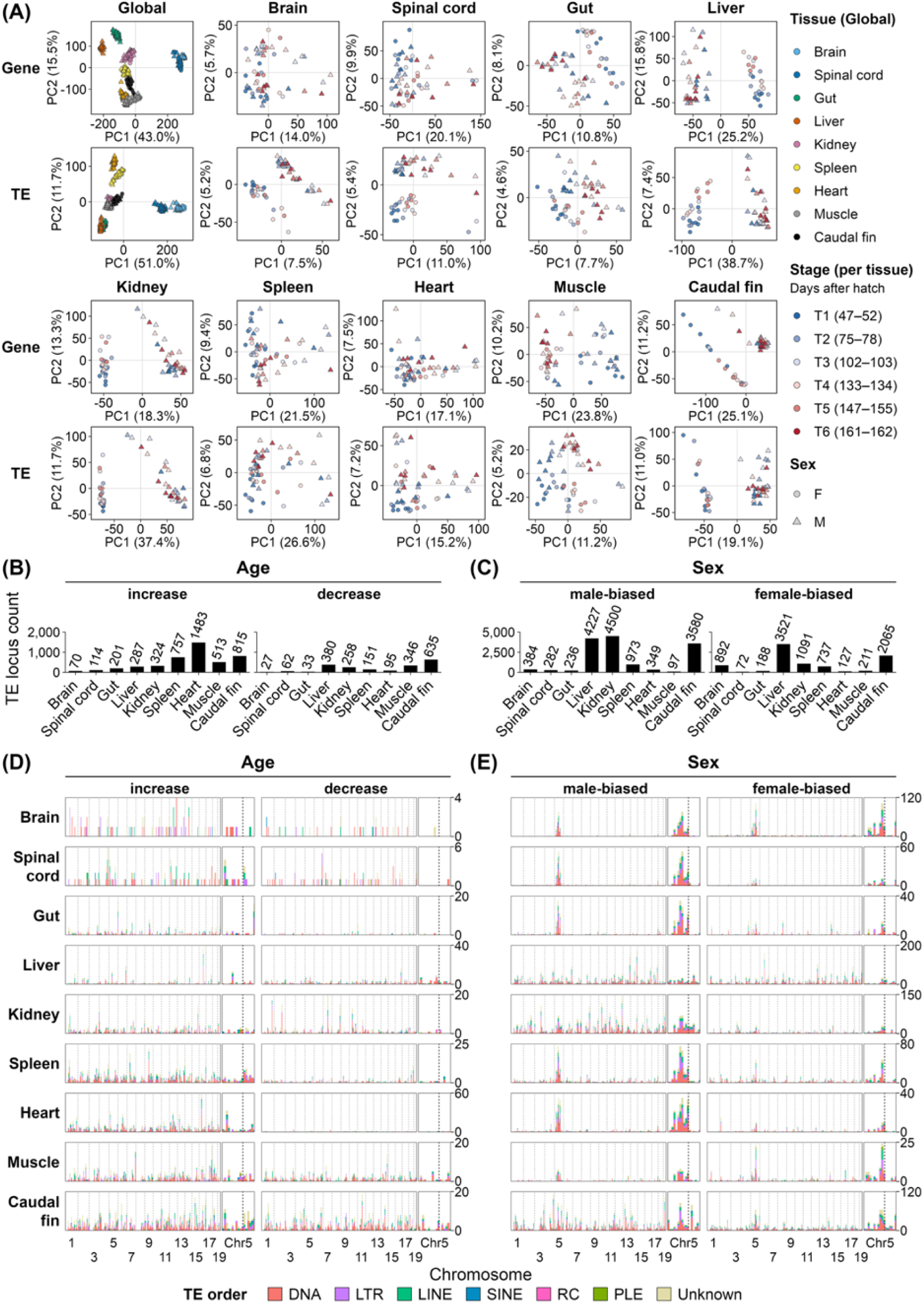
Tissue-dependent patterns of age- and sex-associated TE expression in killifish. (A) Principal component analysis (PCA) of gene expression and TE locus expression across nine tissues and within individual tissues. In the global panels, points are colored by tissue. In the tissue-specific panels, colors indicate age group, and circles and triangles indicate females and males, respectively. Axis labels show the percentage of variance explained by PC1 and PC2. TE expression PCA used loci defined using the Earl Grey-built TE library after tandem repeat masking with ULTRA. (B, C) Numbers of age-associated (B) and sex-associated (C) TE loci across the nine tissues. Counts are shown separately for loci whose expression increased or decreased with age in (B) and for loci with higher expression in males or females in (C). See Methods for the criteria used to identify age- and sex-associated TE loci. (D) Chromosomal distributions of Earl Grey TE loci whose expression increased or decreased with age in the nine tissues. Positive and negative Spearman correlations meeting the age-association criteria in at least one sex define the increase and decrease groups, respectively (Methods). (E) Chromosomal distributions of Earl Grey TE loci showing higher expression in males or females in the same nine tissues. (D, E) Loci were assigned to 5-Mb windows within each chromosome based on their midpoints. Stacked bar heights show the number of loci in each window, and colors indicate TE order. Dotted lines mark chromosome boundaries. In the magnified view of chromosome 5, the *gdf6* locus is indicated by dashed lines. Count scales vary by tissue and between the age- and sex-associated analyses.

Motivated by the TE-based PCA patterns, we tested whether transcript abundance at individual Earl Grey-ULTRA TE loci was associated with age or sex and compared the numbers of age- and sex-associated loci across the nine tissues (Figures 7B and 7C). We assessed monotonic changes in expression with age separately in females and males and retained loci that met the criteria in at least one sex. Sex-associated expression was assessed after adjusting for age group (Methods). Heart and caudal fin showed the largest numbers of age-associated TE loci (Figure 7B). In contrast, relatively few age-associated loci were detected in brain and spinal cord. The numbers of sex-associated TE loci were higher in the liver, kidney, and caudal fin than in the other tissues (Figure 7C). Consistently, these tissues exhibited clear sex-associated separation in the TE-based PCA (Figure 7A).

We next examined the chromosomal distributions of age- and sex-associated TE loci. The numbers of associated TE loci in 5-Mb chromosomal windows were compared across tissues. TE locus counts were summarized separately for age-associated increases and decreases in expression and for male- and female-biased expression (Figures 7D and 7E). Age-associated loci were distributed across all chromosomes, with their chromosomal distributions varying among tissues (Figure 7D). For sex-associated loci, we observed recurrent peaks in the numbers of both male- and female-biased loci on chromosome 5, which harbors the male sex-determining *gdf6* locus^50^, across brain, gut, spleen, heart, and muscle (Figure 7E). The sex-associated peaks on chromosome 5 were consistently observed across all eight TE libraries in both brain and muscle (Figure S6). To examine whether the same TE loci were shared among tissues, we compared the tissue-level sets of age- and sex-associated loci (Figure S7). Sex-associated sets generally showed greater cross-tissue containment than age-associated sets. Together, these results showed that the numbers and chromosomal distributions of age- and sex-associated TE loci varied among tissues, whereas sex-associated loci showed partial cross-tissue sharing and recurrent peaks on chromosome 5.

### Library-dependent detection of age- and sex-associated TE loci

We examined how TE library choice affected the number and overlap of age- and sex-associated TE loci within each tissue, with tandem repeat masking fixed to ULTRA. Because the TE intervals defined using the eight libraries differed in their boundaries and degree of fragmentation, we grouped overlapping intervals from all libraries into a common set of TE meta-loci (Methods). We identified age- and sex-associated TE loci separately for each library, assigned them to the meta-loci, and compared the number and overlap of the meta-loci across libraries (Figures 8A and 8B). Earl Grey, RepeatModeler2, and the RepeatModeler2-built libraries refined with MCHelper or TEtrimmer generally yielded more age- and sex-associated TE meta-loci than the HiTE- and EDTA-built libraries (Figures 8A and B). We then ranked the libraries within each tissue separately for age and sex association (Figures 8C and 8D). For age association, the MCHelper-refined RepeatModeler2-built library ranked first in four tissues and second in one, and Earl Grey ranked first in three tissues and second in four (Figure 8C). For sex association, Earl Grey and RepeatModeler2 showed similar rankings across tissues (Figure 8D). We next examined whether the same TE meta-loci were identified as age- or sex-associated across libraries (Figures 8A and B). The proportion of age-associated meta-loci identified by seven or all eight libraries ranged from 8.2% of the age-associated meta-locus union in brain to 25.7% in heart. The corresponding proportion for sex-associated meta-loci ranged from 21.5% in brain to 44.1% in liver. However, both analyses identified a substantial number of TE meta-loci that met the association criteria only in one library (Figures 8A and 8B). The fraction of age-associated meta-loci detected using only one library was relatively high in the brain (Figure 8A). To examine whether these patterns persisted under more restrictive conditions, we repeated the library comparison using only intergenic TE loci quantified with Telescope’s locus-unique counts (Figure S8). Earl Grey and the RepeatModeler2-derived libraries again yielded relatively large numbers of age- and sex-associated meta-loci, with both shared and library-specific detections remaining evident. Collectively, age- and sex-associated TE meta-locus sets included both loci shared across libraries and loci detected only in one library. Although Earl Grey yielded relatively large sets of associated loci, additional loci identified using other libraries highlighted the value of considering results from multiple TE libraries in locus-level transcriptomic analyses.

**Figure 8.**
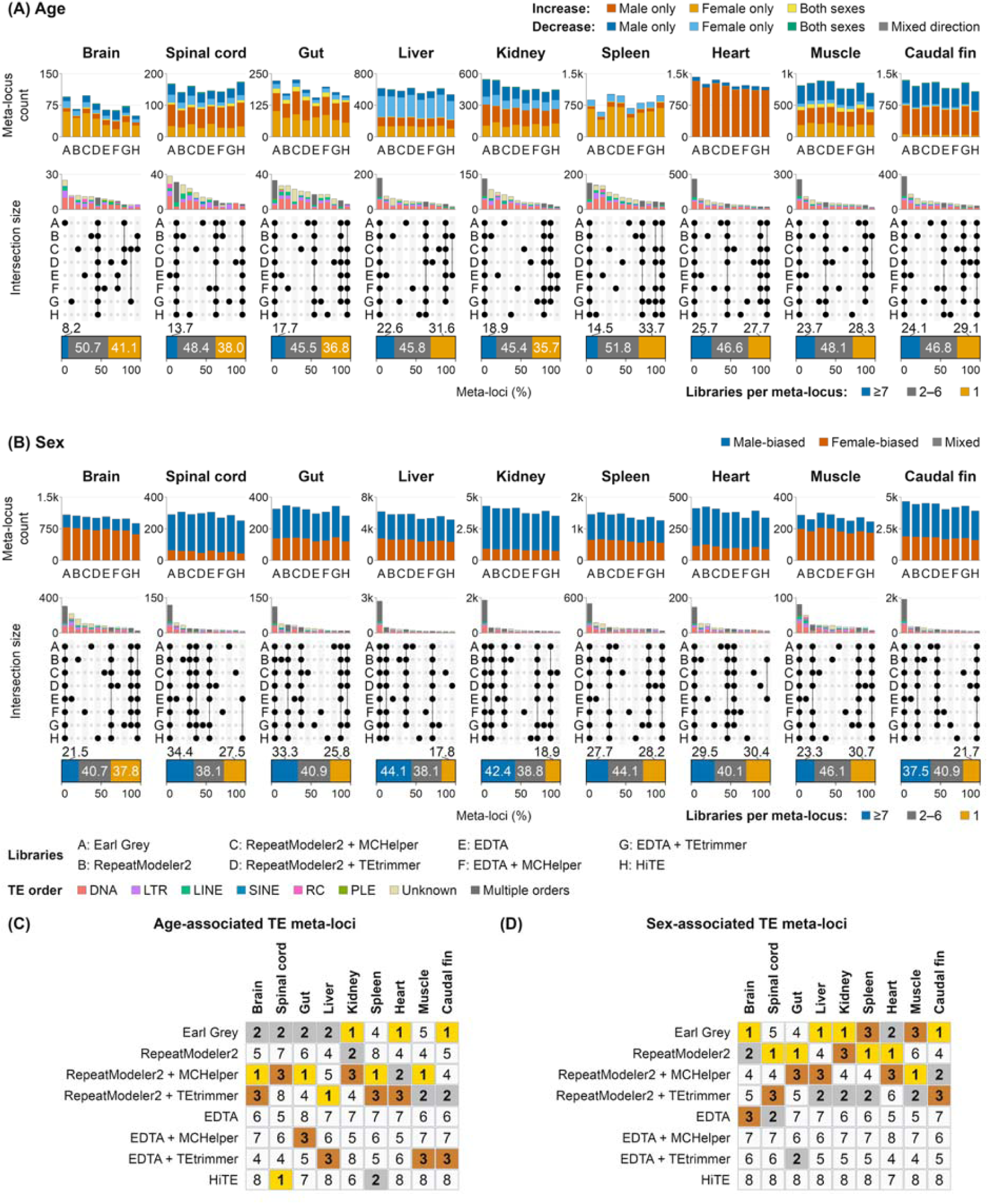
Library-dependent detection of age- and sex-associated TE meta-loci across killifish tissues. All panels are based on combined results from Telescope’s unique-mapping and multi-mapping analyses of both genic and intergenic TE loci. Expression-associated TE loci were mapped to a common set of TE meta-loci, and the union of associated meta-locus sets from the two analyses was used (Methods). (A) Age-associated TE meta-loci in nine tissues. Upper stacked bars show the number of distinct associated meta-loci identified using each TE library. Colors indicate the direction of expression change with age (increase or decrease) and whether the age-association criteria were met in females only, males only, or both sexes. Gray indicates meta-loci with conflicting directions of expression change among contributing results. (B) Sex-differentially expressed TE meta-loci in the same nine tissues. Upper stacked bars show the number of distinct associated meta-loci identified using each TE library. Colors indicate higher expression in females (female-biased) or males (male-biased). Gray indicates meta-loci with conflicting directions of sex bias among contributing results. In (A) and (B), the lower UpSet plots show the 12 largest mutually exclusive intersections among the associated meta-locus sets from the eight libraries. Each bar shows the number of meta-loci identified using exactly the libraries indicated by the connected-dot matrix. The lowest stacked bars summarize all library combinations, including those not shown in the UpSet plots, as percentages of the union of meta-loci across the eight libraries. In both plots, blue denotes meta-loci identified using seven or eight libraries, orange denotes meta-loci identified using a single library, and gray denotes meta-loci identified using two to six libraries. (C, D) Within-tissue rankings of the eight TE library construction workflows based on the expression-associated meta-locus counts in (A) and (B), respectively. Panel (C) shows rankings for age association in at least one sex, and panel (D) shows rankings for differential expression between sexes. Rank 1 indicates the largest number of expression-associated meta-loci, and rank 8 indicates the smallest. Gold, silver, and bronze cells indicate ranks 1, 2, and 3, respectively.

## Discussion

Locus-resolved analysis of repetitive regions depends on their accurate representation in a reference genome assembly. In the present study, we generated a near-complete, chromosome-scale assembly of the African turquoise killifish, designated NfurGRZ-RIMD1. The assembly closely matches the estimated genome size, contains the expected number of chromosome-scale sequences, and recovers satellite-rich regions missing or poorly represented in previous assemblies (Figures 1 and S1). The improved representation of repetitive regions makes NfurGRZ-RIMD1 a useful resource for examining repeat organization and conducting locus-level TE expression analyses. NfurGRZ-RIMD1 is publicly available in the NCBI Genome Data Viewer (https://www.ncbi.nlm.nih.gov/gdv/browser/genome/?id=GCF_043380555.1), together with gene models generated by NCBI’s Eukaryotic Genome Annotation Pipeline (EGAP). The assembly has already been used in several studies, illustrating its value as a reference resource for African turquoise killifish research.^51–54^

Even in the African turquoise killifish, an established vertebrate aging model, the lack of a curated reference set of genomic TE intervals limits evaluation of TE detection accuracy. The need for reliable benchmarking is growing as near-complete genome assemblies become more widely available. To inform the practical selection of repeat-detection workflows, we compared TR detection, TE consensus library construction, and order-level classification methods across near-complete vertebrate assemblies. We then evaluated the genomic coverage and base-pair overlap of TE-derived regions, along with downstream TE expression patterns, across workflows. TR detection parameters altered the apparent TE landscape in genomes containing abundant TRs with long repeat units (Figures 2C and S2). We masked TR regions before TE detection to limit overlapping TE and TR repeat intervals. The TR pre-masking strategy generates repeat tracks with reduced TE-TR overlap for downstream analyses, although studies about repeat evolution may benefit more from retaining TE-only, TR-only, and TE-TR overlapping intervals as distinct categories. Earl Grey-ULTRA yielded the highest combined TE and TR coverage among the 32 combinations tested in each of the three fish species (Figure 6). Locus-resolved analysis of killifish transcriptome data revealed tissue-dependent chromosomal distributions of age- and sex-associated TE loci (Figure 7). Despite broad overlap in TE-derived genomic regions across workflows (Figure 6B), age- and sex-associated TE meta-loci overlapped only partially across workflows (Figure 8). Collectively, our comparison of repeat coverage and expression-associated TE loci reveals distinct detection patterns across workflows without repeat reference intervals.

The Earl Grey and RepeatModeler2 libraries, as well as the RepeatModeler2-derived libraries refined with MCHelper or TEtrimmer, generally yielded more age- and sex-associated TE meta-loci than HiTE and the EDTA libraries across killifish tissues (Figures 8 and S8). A separate benchmark, TE_Bench, found that both Earl Grey and RepeatModeler2 showed higher genome coverage than EDTA.^22^ One possible explanation for the larger number of age- and sex-associated loci is that the Earl Grey library and the other RepeatModeler2-derived libraries provide a broader representation of TE-derived regions for TE locus-level RNA quantification. The differences in TE-derived regions are more likely related to variation in consensus construction and curation because we reclassified all libraries under our majority-vote classification scheme and applied a common TE-derived region detection method (Figure 4A). The relatively large number of expression-associated TE loci identified with RepeatModeler2 without additional curation suggests that the initial consensus library itself influences downstream findings. Curation can further modify the initial library through changes in consensus sequence lengths and library composition. Refinement with MCHelper or TEtrimmer yielded longer consensus sequences in both the RepeatModeler2- and EDTA-built libraries (Figure 5B). Earl Grey also extends consensus sequences, yet the three workflows differ in how they filter consensus sequences and handle redundancy. Earl Grey starts from RepeatModeler2-derived consensus sequences, uses iterative BLAST, Extract, Align, Trim (BEAT) processing to reconstruct and extend consensus sequences, and, unlike MCHelper and TEtrimmer, retains the current consensus when further extension is not supported.^19^ With the default settings of each workflow, the Earl Grey library was used without final-library clustering, whereas the MCHelper- and TEtrimmer-curated libraries were subjected to consensus clustering to reduce redundancy.^17,18^ Thus, curation program differences in sequence extension and retention influence the composition of the final TE library, change the repertoire of TE-derived genomic intervals, and may affect transcript quantification at individual TE loci. Thus, the larger sets of TE loci obtained with Earl Grey and the other RepeatModeler2-derived libraries may stem from both broader starting libraries and subsequent curation of consensus sequences.

Library construction and curation influenced the TE sequence divergence profiles obtained from the same genomes (Figure S5). Previous work showed that incomplete representation of lineage-specific TEs in reference libraries inflates divergence estimates and distorts TE accumulation histories.^55^ The broad sequence divergence distribution observed for the EDTA-built library became more similar to those of the other libraries after refinement with MCHelper or TEtrimmer. In contrast, HiTE, which incorporates its original automated boundary refinement and candidate filtering during library construction^20^, yielded narrow profiles concentrated at low divergence values. These library-dependent sequence divergence patterns persisted when the analysis was restricted to TE-derived regions overlapping genomic intervals identified by all eight libraries (Figure S5B), supporting a contribution of consensus sequence construction to the observed profiles.

Expression at TE loci requires careful interpretation because RNA-seq signals can arise from transcriptional sources other than autonomous TE promoter activity. For intronic TEs, RNA-seq reads can derive from host-gene pre-mRNA or readthrough transcription.^56,57^ In addition, the repetitive nature of TEs can cause RNA-seq reads to map to multiple loci, making read assignment to highly similar TE loci uncertain.^58,59^ Nevertheless, library-dependent patterns in the detection of age- and sex-associated TE loci were generally retained when the analysis was restricted to intergenic TE loci quantified with Telescope’s locus-unique counts (Figure S8). By localizing age- and sex-associated variation in transcript abundance to individual TE loci, our analysis complements previous family-level analyses of TE expression in killifish.^32^ For example, a recurrent peak in the number of sex-associated TE loci on chromosome 5 was observed across several tissues, with the peaks detected using all eight libraries in brain and muscle (Figures 7E and S6). Future studies using long-read RNA sequencing and transcription start site profiling could build on the age- or sex-associated loci to clarify the transcriptional origins and structures of TE-associated transcripts.

In summary, the present study generated a near-complete chromosome-scale assembly and integrated TR detection, TE consensus library construction, TE order classification, TE-derived region detection, and locus-level expression analysis into a single analytical framework. We provide sets of age- or sex-associated TE loci identified across nine tissues using each of the eight libraries (Data S1). The resulting complementary locus sets underscore the value of integrating results from multiple TE libraries for expression analysis and provide a foundation for future studies of TE function and regulation at individual loci in African turquoise killifish.

## Methods

### Animals

The caudal, dorsal, anal, and pectoral fins of the male African turquoise killifish (GRZ-AD) were excised and pooled. Animal handling and sample collection were conducted in accordance with the animal protocol approved by the Institutional Animal Care and Use Committee of Osaka University (RIMD permit #R02-04).

### Genome sequencing and assembly

High molecular weight DNA was extracted from the skeletal muscle using a NucleoBond AXG 500 column (Macherey-Nagel, Düren, Germany). The concentration of the extracted DNA was measured with a Qubit 4 Fluorometer (Thermo Fisher Scientific, MA, USA), whose size distribution was analyzed with a TapeStation 2200 (Agilent Technologies, CA, USA) to ensure high integrity and the length of the main peak in DNA size distribution longer than 50 kb. A SMRT sequence library was constructed based on the official protocol, with a SMRTbell Express Template Prep Kit 2.0 (Pacific Biosciences, CA, USA) and sequenced in a single SMRT cell on a PacBio Revio system (Pacific Biosciences). The sequencing output was processed for generating circular consensus sequences (CCS) to obtain a total of 79.6 Gb of HiFi reads. From these reads, adapter sequences were removed with the program HiFiAdapterFilt.^60^ The obtained HiFi reads were assembled using the program hifiasm v0.19.8^61^ with its default parameters.

### Hi-C library preparation and sequencing

*In situ* Hi-C libraries were prepared using published protocols^62,63^, with the adaptations described below. Caudal, dorsal, anal, and pectoral fins from a single 3.5-month-old GRZ male were briefly wiped with 70% ethanol under anesthesia, excised, and minced. Tissue was dissociated in PBS containing 0.26 U/mL Liberase TM (Merck) for 15 min at room temperature with gentle shaking. Digestion was quenched with fetal calf serum (30–50% of the final volume). After filtration through a 70-µm strainer, cells were collected (350 × *g*, 10 min, room temperature) and resuspended in 2 mL DMEM (approximately 1.2–1.9 × 10 cells per tube). Cells were fixed with 1% formaldehyde for 10 min at room temperature, followed by quenching with 125 mM glycine for 15 min on ice. Cells were collected and washed with ice-cold PBS (350 × *g*, 10 min, 4 °C), then stored as pellets at −80 °C. Frozen pellets were lysed in 25 mL of ice-cold buffer containing 10 mM Tris-HCl (pH 7.5), 10 mM NaCl, 0.2% NP-40, and 1× EDTA-free protease inhibitors. Nuclei were collected and washed with 500 µL of 1.25× NEBuffer 3 (300 × *g*, 10 min, 4 °C), then resuspended in 1 mL of the same buffer and divided into 500-µL aliquots. SDS and Triton X-100 treatments followed Schloissnig et al.^63^, with SDS at 0.5%. Nuclei were pelleted (100 × *g*, 1 min, room temperature) and resuspended in 450 µL of 1× DpnII digestion buffer. Chromatin was digested with DpnII overnight at 37 °C and 950 rpm, followed by biotin fill-in using 50 U of Klenow and ligation as described by Schloissnig et al.^63^. Following proteinase K treatment overnight at 65 °C and RNase A treatment for 2 h at 37 °C, DNA was purified by phenol/chloroform extraction and ethanol precipitation. Biotin removal from unligated ends followed Schloissnig et al.^63^, using 2.5 µg DNA and 3 U of T4 DNA polymerase. DNA was sheared to a peak size of approximately 400 bp using a Covaris S220 (duty factor, 10%; peak incident power, 5; cycles per burst, 200; duration, 70 s). After end repair, DNA was purified using a Monarch DNA purification kit (NEB) and subjected to two-step bead-based size selection. Streptavidin capture was followed by on-bead dA-tailing and Illumina adapter ligation. Capture, adapter ligation, and bead washing followed Schloissnig et al.^63^. Libraries were amplified for eight cycles using NEBNext High-Fidelity 2× PCR Master Mix (NEB), purified with an equal volume of magnetic beads, and quantified using a TapeStation High Sensitivity D1000 assay (Agilent). Sequencing on a NovaSeq X (Illumina) with 150-bp paired-end reads yielded 176 million read pairs.

### Genome scaffolding using Hi-C reads

The obtained Hi-C read pairs were processed using the program Trim Galore! v0.6.8, with the options ‘--phred33 --stringency 2 --quality 30 --length 25 --paired’, and aligned to the obtained HiFi sequence contigs using the program Juicer v1.6.^64^ Based on the read alignment, the HiFi contigs were scaffolded using the program 3d-dna^65^ v201008 with the options ‘-m haploid -i 5000 --editor-repeat-coverage 15 -r 2’, whose output was modified to be consistent with the Hi-C-based chromatin contact profiles. The resultant genome assembly was deposited in NCBI under BioProject <u>PRJNA1162561</u>. For the killifish genome size estimate, we converted the killifish diploid DNA content of 3.11 pg reported by Reichwald et al.^6^ to a haploid genome size using the conversion factor of 0.978 Gb pg⁻¹.^66^

### Gene space completeness

Assembly completeness was evaluated using compleasm v0.2.6^67^ and the actinopterygii_odb10 lineage dataset dated 8 January 2024. Analyses were performed in compleasm’s busco mode (-m busco), with the lineage explicitly specified (-l actinopterygii), and otherwise default analysis parameters. BUSCO’s reference orthologs were classified according to compleasm’s native output. For visualization, interspaced (I) markers were combined with fragmented (F) markers, yielding four categories: single-copy (S), duplicated (D), fragmented (F + I), and missing (M).

### Centromeric repeat profiling

Tandem repeat regions and their consensus repeat unit sequences were identified in the genome assemblies listed in Table S1 using TRASH (v1; Git commit f7d53a0) in R v4.5.1 with default detection and consensus-construction parameters. No predefined repeat-family templates were supplied, and higher-order-repeat analysis was not performed. The consensus monomer sequences in the killifish genomes were compared with centromeric satellite monomer sequences (Nfu-Sat49, Nfu-SatA, and Nfu-SatB), which have been used as FISH probes in a previous study.^6,34^ For sequence comparison, circular shifts were employed, and both forward and reverse complement sequences were included as comparison targets, allowing 1-bp insertions or deletions. We classified a repeat region as a centromeric satellite repeat if sequence identity between each consensus sequence and the repeat region was at least 95%. Sequence identity was calculated as the number of matching bases divided by the length of the FISH probe-derived monomer sequence. To obtain the repeat landscape plot, repetitive structures were visualized for each chromosome using ModDotPlot v0.9.9 in static mode, based on self-dot-plot analysis. The analysis parameters were set with an identity threshold of 86% (-id 86), a k-mer size of 21 (-k 21), a modimizer parameter of 1,000 (-m 1000), a delta of 0.5 (-d 0.5), and a plot resolution of 2,000 (-r 2000).

### Tandem repeat profiles by consensus monomer length

Tandem repeats and their monomer sequences were identified in whole-genome sequences (Table S1) using TRASH as described in the preceding section. Consensus sequence clustering was performed using custom Python scripts. Briefly, repeat abundance was calculated by summing the observed width values of consensus sequences in the TRASH output (consensus.secondary). For each species, the starting position and strand orientation of each consensus sequence were standardized. All circular rotations of the original sequence and its reverse complement were considered, and the lexicographically smallest sequence under the nucleotide order A < C < G < T was selected as the canonical form. Sequences with identical canonical forms were then collapsed into a single entry. The collapsed consensus sequences were further clustered in decreasing order of abundance, defined as the summed lengths of associated TRASH monomers. For each canonicalized consensus sequence, candidate cluster representatives were preselected based on sequence length and 5-mer similarity before alignment-based evaluation. Each query-candidate representative pair was compared using the Python bindings of Edlib v1.3.9.post1, accounting for differences in circular start position and reverse-complement orientation. For each pair, start positions were identified using k-mer matches and semi-global alignments. Sequence identity was defined as one minus the minimum edit distance across the tested alignments divided by the length of the longer sequence in the query-representative pair. Each single-base substitution, insertion, or deletion contributed one to the edit distance. A sequence was assigned to the candidate representative with the highest identity if this value was at least 90%; otherwise, it became the representative of a new cluster. For sequences at least 20 bp long, all length-compatible candidates with a 5-mer-based Jaccard similarity of at least 0.10 were retained for alignment-based evaluation. For sequences shorter than 20 bp, all length-compatible candidates were retained. The abundance of each cluster was calculated as the sum of the observed widths of all final TRASH monomers assigned to that cluster. For representative-length profiles, cluster abundances were grouped by representative consensus sequence length and summed.

### Overlap of TE-derived intervals and TRs

Species-specific *de novo* repeat libraries were generated from whole-genome assemblies of the five selected species (Table S1) using RepeatModeler v2.0.6 in the Dfam TE Tools v1.92 container with default settings. Repeat classification used the Dfam v3.9 Actinopterygii library for the three fish species and the Dfam v3.9 Mammalia library for mouse and human. TE-derived regions in each genome were identified with RepeatMasker v4.1.9 and RMBlast v2.14.1+ (‘RepeatMasker -a -engine rmblast’) using the *de novo* consensus library.

We assigned TE orders to TE-derived intervals based on the label before the first slash in the class/family field of the RepeatMasker output. If the label was DNA, LTR, LINE, SINE, RC, PLE, or Unknown, the TE-derived interval was assigned that label. Other labels were mapped to the target TE labels using common full-name forms as follows: “unclassified” to Unknown; “class II transposon” to DNA; “long interspersed” or “non-LTR retrotransposon” to LINE; “long terminal repeat retrotransposon” to LTR; “short interspersed element” to SINE; “rolling-circle element” or “Helitron” to RC; and “Penelope-like element” to PLE. For labels containing multiple hierarchical levels, the first assignment was used, except that records beginning with LINE/Penelope were assigned to PLE. RepeatMasker categories outside the seven target labels, including “Simple_repeat”, “Low_complexity”, “Satellite”, and RNA-related categories, were excluded. To create mutually exclusive TE class labels per covered base, we generated the genomic interval union of the seven TE labels as follows. We combined all RepeatMasker-identified intervals and split overlapping regions at every interval start and end coordinate, creating non-overlapping segments so that each base pair was counted only once across TE classes. For each resulting segment, we identified all original RepeatMasker-identified regions that covered the segment and assigned it to the TE class of the region with the highest Smith–Waterman score. Exact-score ties were resolved by first preferring intervals not flagged by RepeatMasker as partially overlapping a higher-scoring match, followed by lower percentage divergence, a longer original interval, and earlier appearance in the source file. These priority rules were applied only to overlapping portions. Non-overlapping portions of lower-priority intervals, including those flagged by RepeatMasker as overlapping higher-scoring matches, were retained. Finally, adjacent intervals assigned to the same TE class were merged.

For TR detection, each assembly file was indexed and divided into consecutive, non-overlapping 1-Mb chunks. TRF v4.10.0 was run independently on every chunk at each of six maximum repeat unit lengths (10, 40, 100, 200, 400, and 1,000 bp) using ‘trf <input file> 2 5 7 80 10 50 <maximum_repeat_unit_length< -l 10’. For each species and maximum repeat unit length condition, TRF intervals were sorted by reference sequence and coordinate, and overlapping or directly adjacent intervals were merged.

### Tandem repeat detection, coverage, and overlap analysis

longdust v1.3-r87 was run with its default settings. tantan v51 was run with -s 0.9 -f3, with the maximum tandem repeat unit length specified using -w. ULTRA v1.2.1 was run with -- max_consensus 10 --fdr, with the maximum repeat unit length specified using -p. For TRF v4.10.0, each assembly was split into individual chromosome-scale sequences, which were processed in parallel using the parameters 2 5 7 80 10 50, followed by the species-specific maximum repeat unit length and the option -l 10. For tantan, ULTRA, and TRF, the maximum repeat unit length was set to 400 bp for killifish, 300 bp for zebrafish, 600 bp for medaka, 300 bp for mouse, and 400 bp for human. Unless otherwise specified, default settings were used for all program options. For each species and program, identified tandem repeat intervals were sorted by sequence name, start, and end and merged with bedtools v2.31.1 merge using its default distance rule. Within each species and program, merged intervals of the same length were grouped, and their total coverage was calculated as the interval length multiplied by the number of intervals. Cumulative coverage was then calculated by summing coverage across all interval lengths up to each threshold.

For mutually exclusive base-pair intersections, the merged interval sets for each program and species were analyzed with bedtools v2.31.1 multiinter. Each genomic interval identified by at least one program was grouped according to the set of programs that identified the interval, and interval lengths were summed for each program combination. Within each species, program combinations were ranked from highest to lowest exclusive coverage.

For pairwise comparisons between programs, the total coverage for each program was calculated by summing the lengths of the merged intervals. For each of the six pairs of programs within each species, the total base-pair overlap was calculated using bedtools jaccard with the minimum overlap of 1 bp and without imposing an additional fractional or reciprocal-overlap threshold. Directional containment was then calculated by dividing the shared base-pair overlap by the total size of the coverage generated by the row program.

### TE consensus sequences

#### Dfam

Dfam reference libraries were retrieved from Dfam v3.9 FamDB using Dfam TE Tools v1.95. Each library included curated TE consensus sequences associated with the target species and its ancestral taxa. TE order class names were retained in the sequence headers.

#### Earl Grey

Earl Grey v6.1.1 was run on unmasked assemblies without prior RepeatMasker masking against known repeats. The internal RepeatModeler v2.0.6 step was performed without LTR structural analysis. Optional consensus library clustering and HELITRON detection were disabled.

#### RepeatModeler2

RepeatModeler2 v2.0.6 was run in the Dfam TE Tools v1.92 container, using RMBlast v2.14.1+ as the sequence search engine. The optional LTR structural-discovery pipeline was disabled in all runs. The consensus sequences generated were used as input to MCHelper and TEtrimmer.

#### EDTA

Libraries were generated with EDTA v2.2.2. We enabled RepeatModeler-based detection to identify additional repeats during library construction. Protein-coding sequences from killifish and zebrafish, used to filter spurious TE consensus sequences, were retrieved from the corresponding RefSeq genome sequences GCF_043380555.1 and GCF_049306965.1, respectively. The medaka protein-coding sequences were retrieved from the previous study.^36^ The resulting nonredundant TE consensus sequences were used as inputs to MCHelper and TEtrimmer.

#### MCHelper

Each EDTA-built and RepeatModeler2-built library was curated separately with MCHelper (Git commit v1.7.0-9-g984a669). MCHelper was run with all modules enabled in fully automatic mode (-r A -a F). The HMM profile file supplied was the Pfam 35.0 database.

#### TEtrimmer

Each EDTA-built and RepeatModeler2-built library was curated separately with TEtrimmer v1.5.4 using the --classify_all option.

#### HiTE

HiTE v3.3.2 was run separately on each assembly in a Singularity container, with the input specified as a non-plant genome.

The resulting consensus TE libraries were screened with ULTRA v1.2.1 using a maximum detectable repeat unit length of 1,000 bp and a minimum of two repeat units. For each consensus, ULTRA-detected repeat intervals were merged, and TE consensus sequences were removed if tandem repeats detected by ULTRA covered ≥80% of the consensus sequence length.

### Classification of TE consensus sequences

TE consensus sequences from Dfam and *de novo* libraries were classified using RepeatClassifier v2.0.7, Terrier v0.3.3, CREATE (Git commit 4f686b6), NeuralTE v1.0.1-17-g18e99a9, and TEclass2 (Git commit 612470e). For RepeatClassifier-based classification of the *de novo* libraries from killifish, zebrafish, and medaka, a reference library was prepared from Dfam v3.9 by extracting both curated and uncurated families associated with Actinopterygii and its descendant and ancestral taxa. RepeatClassifier was run in the Dfam TE Tools v1.95 container. Terrier was run with the options --min-length 20 --max-length 15000. CREATE was run with its pretrained hierarchical model and default inference settings. NeuralTE was run using the pretrained NeuralTE-TSDs model with target site duplication (TSD) features enabled. All input consensus sequences were classified with NeuralTE, including those for which no TSD was detected. TEclass2 was run in classification mode using the clust_cats_16 model configuration. TEclass2 predictions with a maximum class probability below 0.8 were assigned “Unknown” during output parsing.

TE order labels were harmonized across classifiers before comparison. First, blank labels were set to “Unknown”. We then converted TIR and Helitron to DNA and RC, respectively, and grouped rRNA, snRNA, tRNA, scRNA, and RNA as Structural_RNA. DIRS labels from CREATE were converted to LTR. For the classifier evaluation with the Dfam reference libraries, only consensus sequences with reference order labels of DNA, LTR, LINE, SINE, RC, or PLE were retained.

For classifier comparisons using *de novo* TE consensus sequences, the pairwise exact-match rate for each classifier pair was calculated as n_same_teorder_/n_common_teid_, where n_common_teid_ is the number of TE consensus sequence IDs present in both classifier outputs and n_same_teorder_ is the number of TE IDs for which the two classifiers assigned identical TE order labels.

### Consensus TE order labels

We evaluated all six two-classifier combinations, all four three-classifier combinations, and one four-classifier combination under strict or majority-vote rules. For strict consensus order label classification, we retained a TE order only when all constituent classifiers predicted the same order label; otherwise, the consensus was set to “Unknown”. For each three-classifier combination under the majority-vote rule, the TE order supported by at least two classifiers was assigned to the consensus, whereas sequences for which all three classifiers predicted different orders were labeled “Unknown”. “Unknown” was treated as a regular category in the majority vote. For the evaluation of Dfam reference consensus sequences in Figure 4D, sequences assigned “Unknown” were excluded from the label agreement calculation. Label agreement was calculated as the percentage of classifier-predicted sequences whose assigned order matched the Dfam reference. In contrast, the percentage of “Unknown” assignments was calculated using all evaluated sequences as the denominator.

### Refinement of the NfurGRZ-RIMD1 gene models

We refined killifish gene models with paired-end RNA-seq datasets from the multi-tissue aging atlas of African turquoise killifish (GRZ strain).^49^ For each sample of the 14 tissues (bone, brain, caudal fin, fat, gut, heart, kidney, liver, muscle, ovary, retina, spinal cord, spleen, and testis), we processed the paired-end read files with fastp v1.3.2 using default settings. A STAR v2.7.11b index was built from the NfurGRZ-RIMD1 assembly and its matched NCBI RefSeq gene model (GCF_043380555.1-RS_2024_12), using -- sjdbOverhang 149. Trimmed read pairs were aligned to the index with STAR v2.7.11b using --outSAMstrandField intronMotif, and coordinate-sorted BAM files were generated for transcript assembly. Other alignment parameters were left at their defaults.

For reference-guided transcript assembly, each alignment file was assembled independently with StringTie v2.2.3. No additional StringTie or strand-specific options were supplied. The sample-level transcript models were first merged separately within each of the 14 tissues using StringTie --merge with otherwise default settings. The resulting 14 tissue-level transcript models were then merged with the same command to generate the all-tissue transcript models. Novel transcripts that did not overlap with the original gene models were removed, and the resulting transcript models were used to filter exon-overlapping TE loci as described in the following section.

### TE and TR genomic landscapes

Nucleotide bases identified as TRs by ULTRA, TRF, tantan, or longdust were hard-masked with “X”. Each hard-masked assembly was screened independently against each of the eight *de novo* TE libraries, which had been filtered to remove TR-containing consensus sequences and classified by the three-classifier majority voting as described in the preceding sections. RepeatMasker v4.2.2 was run in the Dfam TE Tools v1.95 container using RMBlast as the search engine, with alignment output enabled and masking of low-complexity and small RNA sequences disabled.

We next incorporated TR intervals generated by the four TR detection methods into the corresponding RepeatMasker outputs. For each TR method, overlapping TR intervals were first merged using bedtools merge. The merged TR intervals were then combined with the TE-derived regions that RepeatMasker identified. To join fragmented TE intervals, RepeatMasker fragments were grouped by the RepeatMasker ID (column 15), chromosome or scaffold, and the name of the matched TE consensus sequence. Fragments in each group were joined into a single interval extending from the minimum query start to the maximum query end. When two processed intervals overlapped by at least 50% of the length of the shorter interval, the shorter interval was marked for removal. All overlapping pairs were evaluated independently, including pairs involving intervals already marked for removal. If their lengths were equal, the interval appearing first in the input file was retained. All marked intervals were removed after the pairwise comparisons. Any remaining repeat intervals overlapping an exon by at least 1 bp were then removed irrespective of sequence strands. Exon coordinates were taken from the refined gene model (see the section “Refinement of the NfurGRZ-RIMD1 gene models”) for killifish, the RefSeq gene model of GCF_049306965.1 for zebrafish, and the gene model reported in the previous study^36^ for medaka.

For TE locus expression analysis, we used TE-derived regions identified by RepeatMasker after hard-masking ULTRA-detected TRs. Tandem repeat intervals were not added to the RepeatMasker output. RepeatMasker fragments were grouped and joined as described in the preceding paragraph. Both processed and unprocessed intervals shorter than 100 bp were discarded. The remaining intervals were further filtered using the same overlap and exon exclusion criteria described above. The resulting intervals were used as reference loci for TE-Seq analysis, with a unique gene_id assigned to each locus.

### Quantification of TE and TR coverage

Genome-wide repeat coverage was quantified using intervals generated by merging TE and TR regions and applying the filtering described in the preceding section. To avoid counting overlapping bases more than once, we calculated coverage separately for each chromosome by splitting overlapping repeat regions at all interval start and end coordinates to generate non-overlapping segments. Regions covered by a single interval were assigned entirely to that interval. For regions covered by exactly two intervals, half of the region length was assigned to each interval, regardless of repeat category, interval length, or strand. Fractional base-pair values were retained when the overlapping region had an odd number of base pairs. The adjusted coverage was then summed by repeat category. The unmasked genome fraction was calculated as the proportion of bases not covered by any of the repeat intervals.

### TE sequence divergence

TE sequence divergence was analyzed from RepeatMasker v4.2.2 alignments generated by screening ULTRA hard-masked genome assemblies against each of the eight *de novo* TE consensus libraries. We used calcDivergenceFromAlign.pl to summarize CpG-adjusted Kimura two-parameter divergence between genomic TE copies and their assigned consensus sequences. The resulting files contain, for each species-library combination, the length-weighted mean divergence, total aligned length (absLen), and aligned length used for divergence calculation (wellCharLen) for each consensus sequence. Records with missing divergence values or absLen = 0 were excluded. Negative divergence values were set to zero. Records were retained only if wellCharLen/absLen ≥ 0.5 and wellCharLen > 30 bp. For each retained record, genomic abundance was calculated as (wellCharLen) / (total assembly length). Records were grouped by length-weighted mean divergence into 2-percentage-point bins: [0, 2), [2, 4), …, [24, 26), with values ≥26% assigned to a final “26+” bin. Genomic abundance was then summed within each species-library combination and divergence bin (Figure S5A).

For Figure S5B, we selected genomic intervals identified as TE-derived by all eight libraries, based on the genome-wide overlap analysis shown in Figure 6B. Adjacent intervals were merged to define a continuous interval set for each species. For each original RepeatMasker alignment, we counted the query bases overlapping these intervals on the same chromosome, counting each base only once and ignoring strand. An alignment was retained if the total overlap covered at least 50% of its genomic query span. Retained alignments were used in full, including gaps and sequence outside the shared intervals. The retained alignments were then processed with calcDivergenceFromAlign.pl and summarized using the same filtering, normalization, and binning procedures described above.

### Gene and TE locus transcript quantification

We retrieved paired-end RNA-seq data from African turquoise killifish of the GRZ strain from the multi-tissue aging atlas.^49^ The tissues used were bone, brain, retina, fat, gut, heart, kidney, liver, muscle, ovary, caudal fin (source tissue name: skin), spinal cord, spleen, and testis. We omitted five tissues from the subsequent analysis because they were female- or male-only organs (testis and ovary), had fewer biological replicates (retina and ovary), or had low-quality RNA (retina, bone, and fat), according to the original study. Collection stages T1-T6 correspond to 47-52, 75-78, 102-103, 133-134, 147-155, and 161-162 days after hatching, respectively. RNA-seq data were processed using a TE-Seq workflow (Git commit 9bcc4f1) with Snakemake v9.16.3 and the maxfieldkelsey/te-seq:v1.3 container executed through Singularity. Reads were processed with fastp v0.24.1 using paired-end adapter detection and aligned to the NfurGRZ-RIMD1 assembly with STAR v2.7.11b. STAR allowed up to 1,000 multi-mapping alignments per read (--winAnchorMultimapNma× 1000 and -- outFilterMultimapNma× 1000) and emitted intron-motif strand tags.

For gene transcript quantification, genes were quantified against the RefSeq gene model (GCF_043380555.1-RS_2024_12) with featureCounts v2.1.1. Paired-end primary alignments were counted with -p -s 0 -O -M -t exon --minOverlap 25 --fraction --primary. Only primary alignments were counted. Reads overlapping multiple genes contributed fractional counts. For TE locus transcript quantification, the processed TE intervals as described in the section “TE and TR genomic landscapes” were converted to reference loci for TE-Seq analysis, with a unique gene_id assigned to each locus. TE reads were assigned with Telescope v1.0.3+49.g2832514 using telescope assign --attribute gene_id --stranded_mode None --reassign_mode average. We analyzed both Telescope’s locus-unique counts and its multimapping-inclusive counts.

For each tissue, gene and Telescope counts were combined, missing TE entries were set to zero, and fractional counts were rounded to the nearest integer before construction of a DESeq2 dataset. DESeq2 v1.38.0 under R v4.2.3 was used to estimate one size factor per sample, with only gene rows specified as control. The size factor was calculated solely on featureCounts-derived gene counts shared between the unique- and multimapping modes. The resulting size factors were applied to both gene and TE counts in both mapping modes. No batch variable was included in the model, and no additional batch correction was applied to the normalized counts.

### Principal component analysis of gene and TE locus expression

#### Global gene PCA

Gene counts generated by featureCounts were normalized using size factors estimated from Telescope-unique gene counts, as described above. We combined the count tables normalized independently for each tissue. A gene was retained if its normalized count was at least 5 in at least 3 samples from any one tissue. Counts were transformed as log2(normalized count + 1). For each retained gene, sample variance was calculated across all RNA-seq libraries from the nine selected tissues, pooling all represented age groups and sexes. Each RNA-seq sample contributed one observation to variance ranking and PCA. Genes were ranked by variance, and the top 5,000 genes were selected with ties broken by ascending gene ID. PCA was performed on the resulting sample-by-gene matrix after centering each gene on its mean across all RNA-seq libraries, without unit-variance scaling. PCA did not apply VST, rlog, CPM/TPM conversion, quantile normalization, or any additional library-size normalization. Global gene PCA was performed in R v4.5.1, using data.table v1.18.2.1 for data handling, and matrixStats v1.5.0 for variance calculation.

#### Global TE locus PCA

Telescope’s multimapping-inclusive transcript abundance estimates were used for TE loci defined by the Earl Grey-ULTRA workflow. A TE locus was eligible when its normalized count was at least 5 in at least 3 samples within any one of the tissues. Values were transformed as log2(normalized count + 1). Sample variance was calculated for each eligible locus across all libraries, zero-variance loci were excluded, and the 5,000 loci with the largest variance were selected with ties broken by ascending locus ID. These selected loci were analyzed using the same centered, unscaled PCA procedure and R environment as used for the gene PCA across all tissues.

#### Per-tissue PCA

Telescope’s multimapping-inclusive transcript abundance estimates were used for TE loci defined by the Earl Grey-ULTRA workflow. Genes and TE loci were retained if their normalized counts were at least 5 in at least 3 samples within any one of the 14 tissues in the original dataset. For each of the nine displayed tissues, sample variance was calculated across libraries from that tissue using log2(normalized count + 1) values. The 5,000 most variable features were selected separately for genes and TE loci. PCA was performed after centering each selected feature on its within-tissue mean, without unit-variance scaling.

### Age-associated TE loci

Age-associated expression analysis was performed in Python v3.13.0 using NumPy, pandas, and SciPy. The T1-T6 size-factor-normalized count tables were used without recalculating size factors. Within each tissue, repeat library, Telescope mode, and sex, a locus was defined as testable when its normalized count was greater than 1 in at least half of the samples of that sex, with the required number rounded up for odd sample sizes. Spearman rank correlation was calculated between normalized count and the animal’s exact age in days separately for females and males. Two-sided *P* values for Spearman’s ρ were approximated using a t distribution with *n* - 2 degrees of freedom, where *n* denotes the number of samples included in the test. The resulting *P* values were adjusted using the Benjamini–Hochberg procedure separately within each combination of sex, tissue, repeat library, and Telescope mode. For each sex, we classified a locus as age-associated if it had an absolute Spearman’s ρ greater than 0.50 and a Benjamini–Hochberg-adjusted *P* value below 0.10. Age-association status was evaluated independently for females and males.

### Sex-associated TE loci

Sex-differential expression analysis was performed in R v4.2.3 using DESeq2 v1.38.0 and apeglm v1.20.0. All T6 samples were excluded before normalization and model fitting because this stage included males only. For each tissue-by-library-by-Telescope-mode dataset, size factors were re-estimated from rounded gene transcript counts using only T1-T5 samples and then applied to the corresponding raw integer TE transcript counts. Expression filtering was performed separately for each sex. A locus passed the filter for a given sex if the normalized count of a locus exceeded 1 in at least half of the samples of that sex, with the required number of samples rounded up when the sample size was odd. Loci passing the filter in either sex were modeled in DESeq2 with stage as a factor with levels T1-T5 and the design formula ∼ stage + sex. The tested DESeq2 coefficient represented the log2 fold change in expression in males relative to females. No stage-by-sex interaction term was included. DESeq2 was run using a two-sided Wald test, parametric dispersion fitting, and no beta prior, with automatic replacement of outlier counts disabled. Independent filtering was disabled, and nominal *P* values were adjusted using the Benjamini–Hochberg procedure separately within each expression-filtered tissue-by-library-by-mode dataset. Log2 fold changes were shrunk with apeglm using type = “apeglm” and apeMethod = “nbinomC”. A locus was classified as a sex-differentially expressed locus if it had a Benjamini– Hochberg-adjusted *P* value < 0.05, an absolute apeglm-shrunken log2 fold change ≥ 1, successful convergence of the DESeq2 coefficient estimates, and was not flagged as a Cook’s-distance outlier by DESeq2.

### Chromosomal distributions of age- and sex-associated TE loci

TE loci meeting the age- or sex-association criteria described above were used to summarize chromosomal distributions of TE loci. For each tissue and TE library, expression-associated loci identified using Telescope’s unique-mapping and multi-mapping analyses were combined, including both genic and intergenic loci. For age association, loci meeting the criteria in at least one sex were included. Locus counts were based on the library-specific source TE loci used for transcript quantification. Each source locus was assigned to a 5-Mb bin within its chromosome based on the midpoint of the TE locus interval. Within each plotted group, repeated records sharing the same source TE locus ID in both the unique-mapping and multi-mapping analyses were counted only once.

### TE meta-locus counts

Before TE workflow comparison, all TE-Seq-processed TE intervals from the eight libraries were pooled to define a fixed set of canonical TE meta-loci. We sorted the intervals and grouped those connected by overlaps of at least 1 bp into meta-loci. Intervals were connected either directly or through a chain of overlapping intervals. Adjacent intervals were not merged unless they overlapped. Each source locus was assigned to exactly one meta-locus ID, and all meta-loci were retained, including those represented in only one library. For Figure 8, we mapped age- and sex-associated TE loci identified using Telescope’s multi-mapping and unique-mapping modes to the canonical meta-loci. We then took the union of the associated meta-locus sets from the two modes. For Figure S8, we restricted the analysis to intergenic TE loci and used Telescope’s locus-unique counts. We classified age-associated meta-loci as female-only, male-only, or supported in both sexes based on whether at least one contributing source locus met the corresponding sex-specific criterion for age association. Support in both sexes required at least one qualifying result for females and at least one for males within the same meta-locus. These results did not need to come from the same source record or mapping mode. When the contributing results indicated both increases and decreases in expression with age, the meta-locus was labeled “Mixed”. For differential expression between sexes, the direction of each contributing result was determined from the sign of the shrunken log2 fold change for males relative to females. Meta-loci with conflicting directions among contributing results were also labeled “Mixed”.

### Generative AI assistance in data analysis and visualization

ChatGPT (OpenAI; using the latest available model versions at the time of use; May 2025– October 2026) and Codex CLI (OpenAI; regularly updated during the study period; version 0.160.0 at the time of manuscript preparation, using the latest available model at the time of use) were used to assist with bioinformatic analyses, the development and revision of analysis and visualization code, and advice on statistical methods. AI tools were also used to check the analysis and code. Plots were generated from the outputs of the analyses described above using R v4.5.1 graphics, ggplot2 v4.0.2, patchwork v1.3.2, grid v4.5.1, gtable v0.3.6, and scales v1.4.0. We reviewed all resulting plots and made the final scientific interpretations. Availability of the analysis and visualization scripts and figure source data is described in the “Data and code availability” section.

## Data and code availability

The NfurGRZ-RIMD1 genome assembly has been deposited in NCBI under BioProject PRJNA1162561 and is available under RefSeq accession GCF_043380555.1. The PacBio HiFi sequence reads and Hi-C read pairs will be available in the NCBI Sequence Read Archive (SRA) under run accessions SRR38762123 and SRR38762414, respectively. The TE consensus libraries, genomic repeat intervals, refined gene model, expression matrices, statistical results, and source data underlying the figures will be available on Zenodo. Custom analysis scripts will be available on GitHub. Large datasets not deposited on Zenodo are available from the lead contact upon request.

## Acknowledgments

The authors thank Akifumi Yao, Takuma Niida, and Yawako W. Kawaguchi for valuable comments on the manuscript and Hatsune Makino-Itou for technical assistance in whole-genome sequencing. Computations were partially performed on the NIG supercomputer at the ROIS National Institute of Genetics.

## Author contributions

Conceptualization, Y.O. and S.K.; methodology, Y.O., A.K., K.A., and S.K.; investigation, Y.O., A.K., K.A., and S.K.; formal analysis, Y.O.; data curation, Y.O. and S.K.; resources, K.A.; visualization, Y.O.; software, Y.O.; writing – original draft, Y.O.; writing – review and editing, Y.O., A.K., and S.K.; supervision, T.I. and S.K.; funding acquisition, Y.O., A.K., T.I., and S.K.

## Supplemental information

Figures S1-S8, Table S1, and Data S1.

## Declaration of generative AI and AI-assisted technologies in the manuscript preparation process

During the preparation of this work, the authors used ChatGPT and Codex (OpenAI) to assist with manuscript drafting and revision, and literature searches. Details of AI assistance in data analysis and visualization are provided in the Methods section. The authors reviewed and edited the manuscript as needed and reviewed all figures. They take full responsibility for the content of the published article.

