## Supplemental information for "Near-complete genome assembly of the African turquoise killifish enables locus-level profiling of transposable element expression"

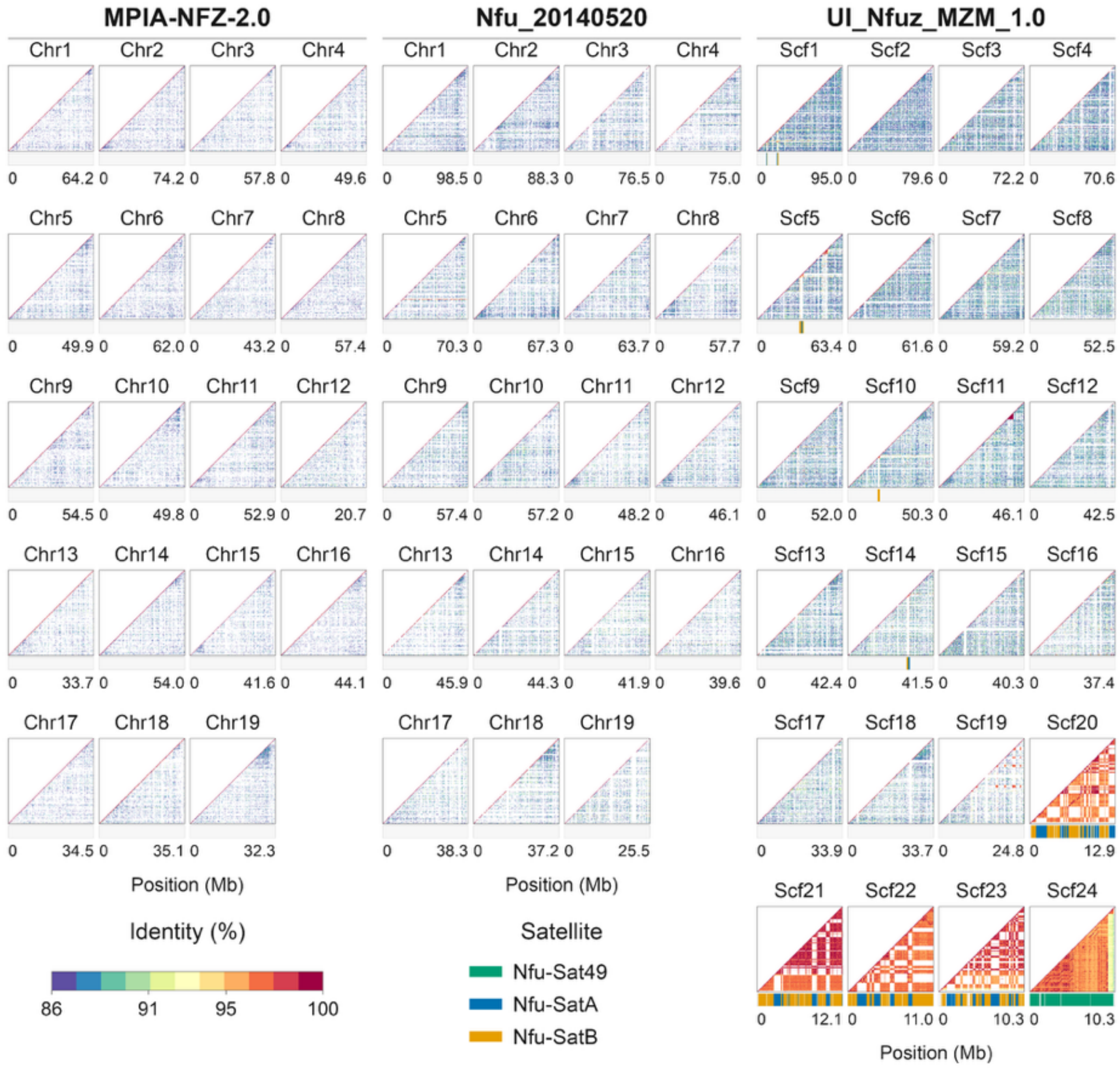


**Figure S1. Distribution of repetitive regions in previously published killifish genome assemblies.** Plot format and color coding are as in Figure 1B.

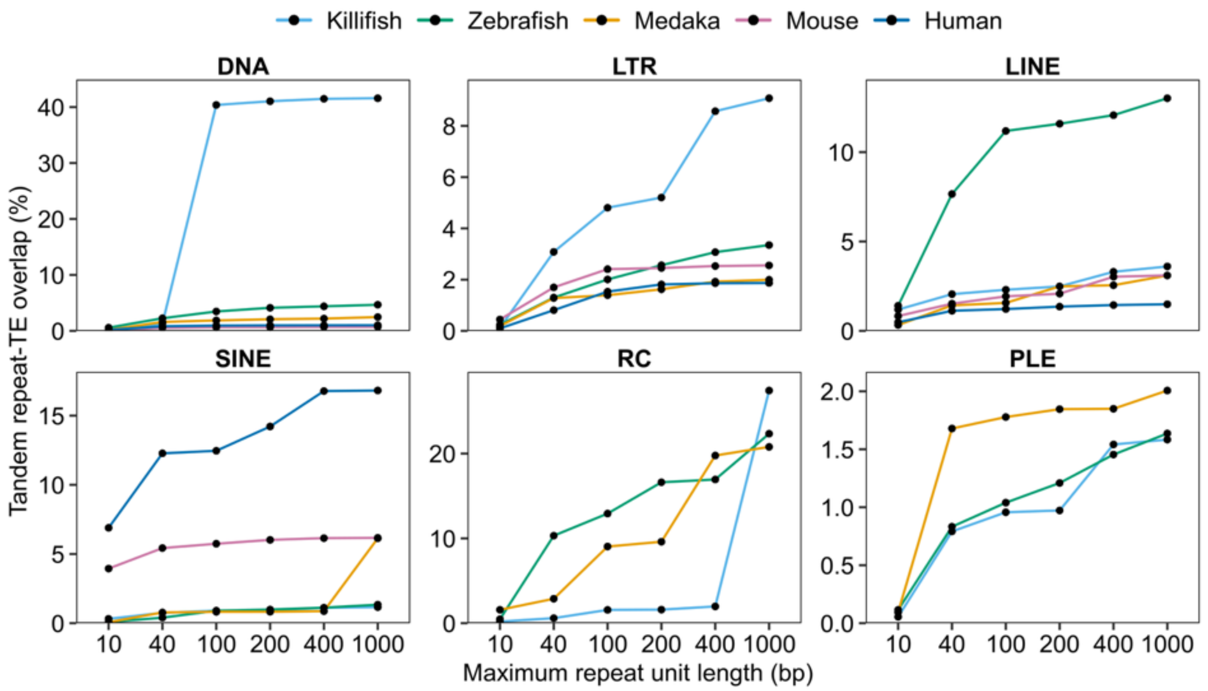


**Figure S2. TE order–specific overlap with tandem repeats across TRF parameter settings.**

Each panel shows the percentage of bases in the indicated TE order that overlap TRF intervals. Line colors indicate species, as in Figure 2C.

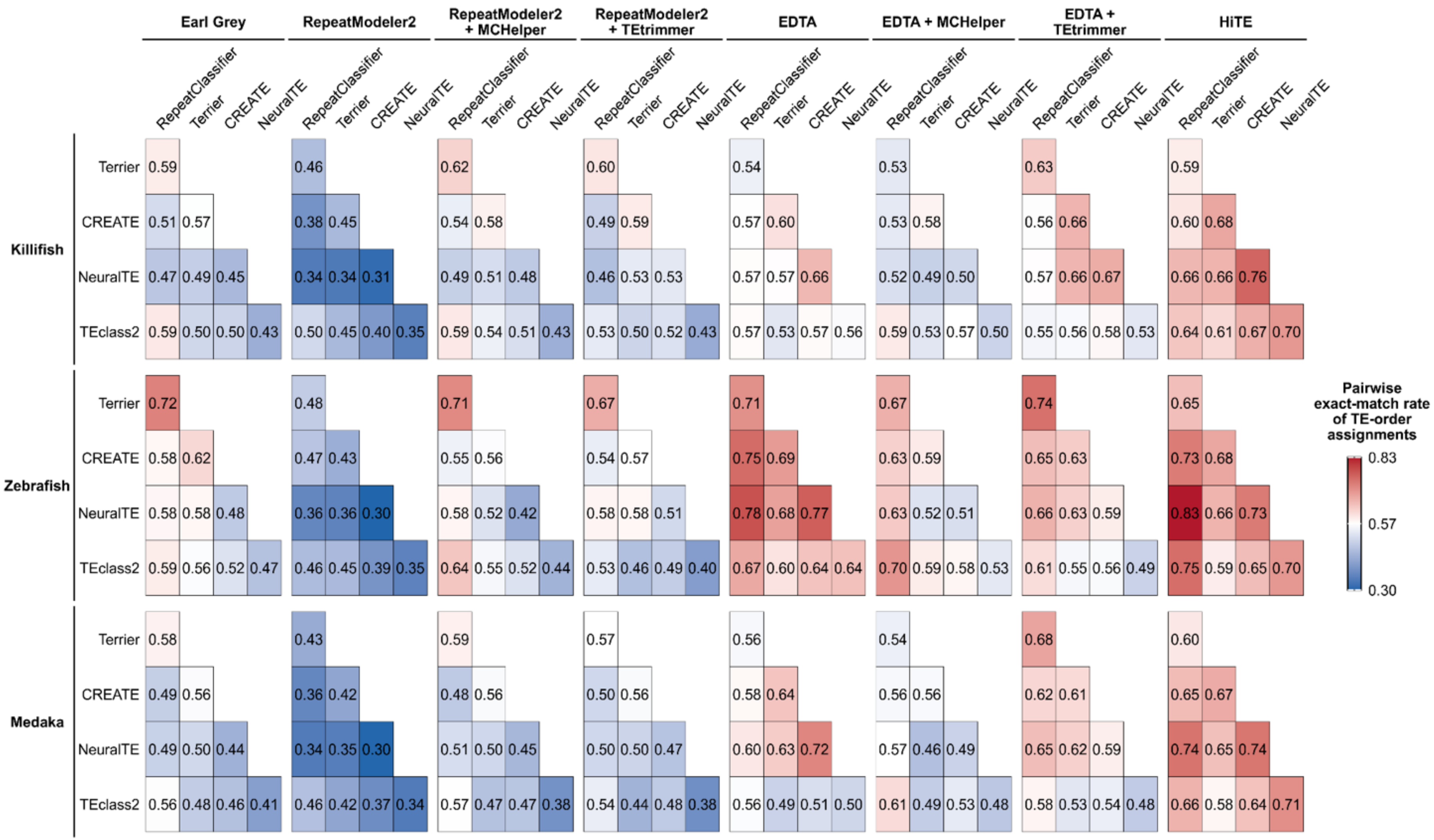

**Figure S3. Pairwise exact-match rates of TE order assignments across de novo libraries.**

Related to Figure 4C. For each species and de novo TE consensus library, the pairwise exact-match rate is the fraction of sequences classified by both classifiers that were assigned the same TE order.

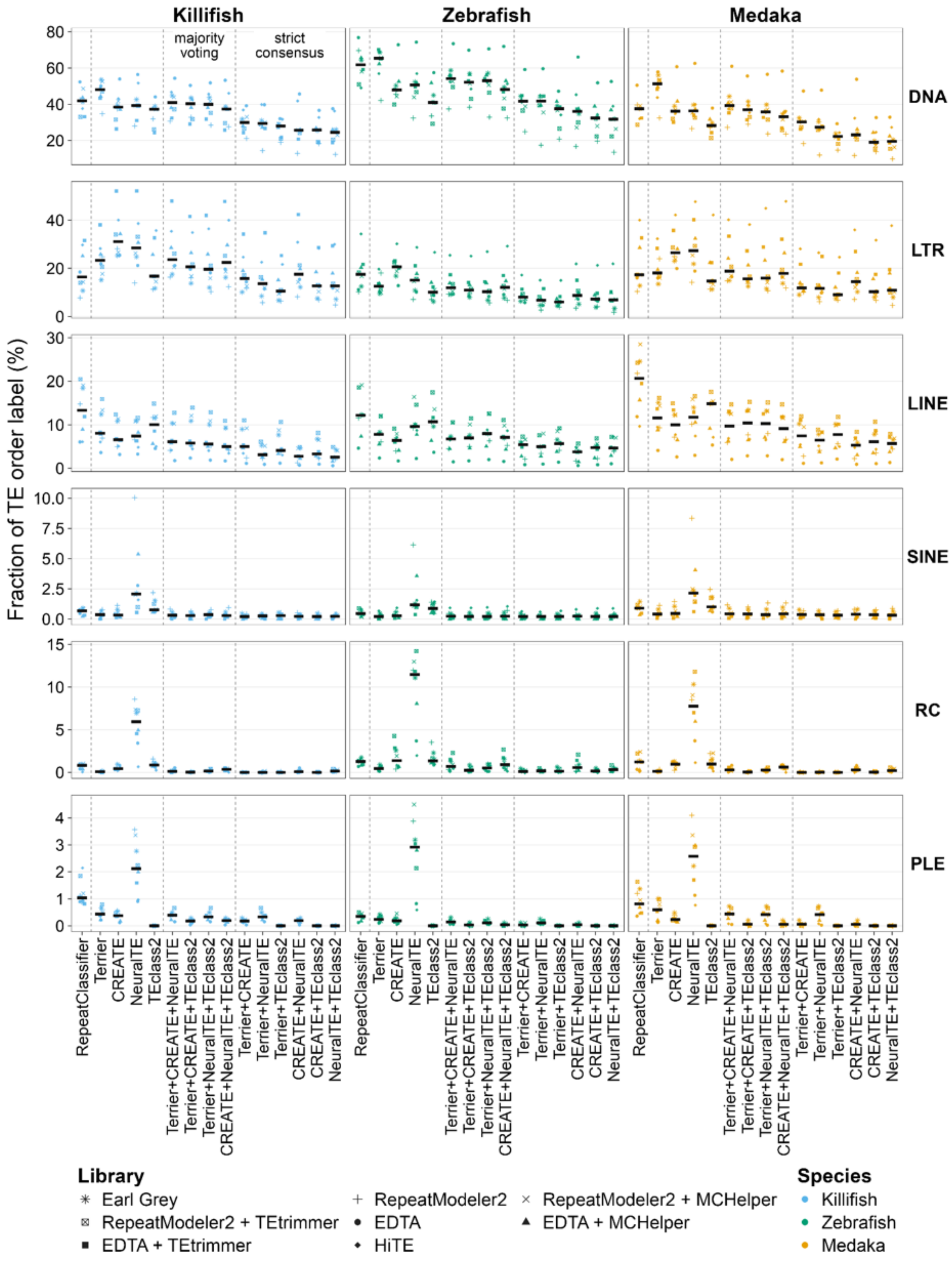


**Figure S4. TE order composition of de novo libraries across classification strategies.**

Related to Figure 4E. Rows represent TE orders, and columns represent species. Each point shows the percentage of TE consensus sequences in one library assigned to the indicated TE order under the classification strategy shown on the x-axis. Black horizontal bars indicate the median across the eight TE libraries for each species, TE order, and classification strategy. Dashed vertical lines separate the groups for RepeatClassifier, individual machine learning classifiers, majority voting, and strict consensus.

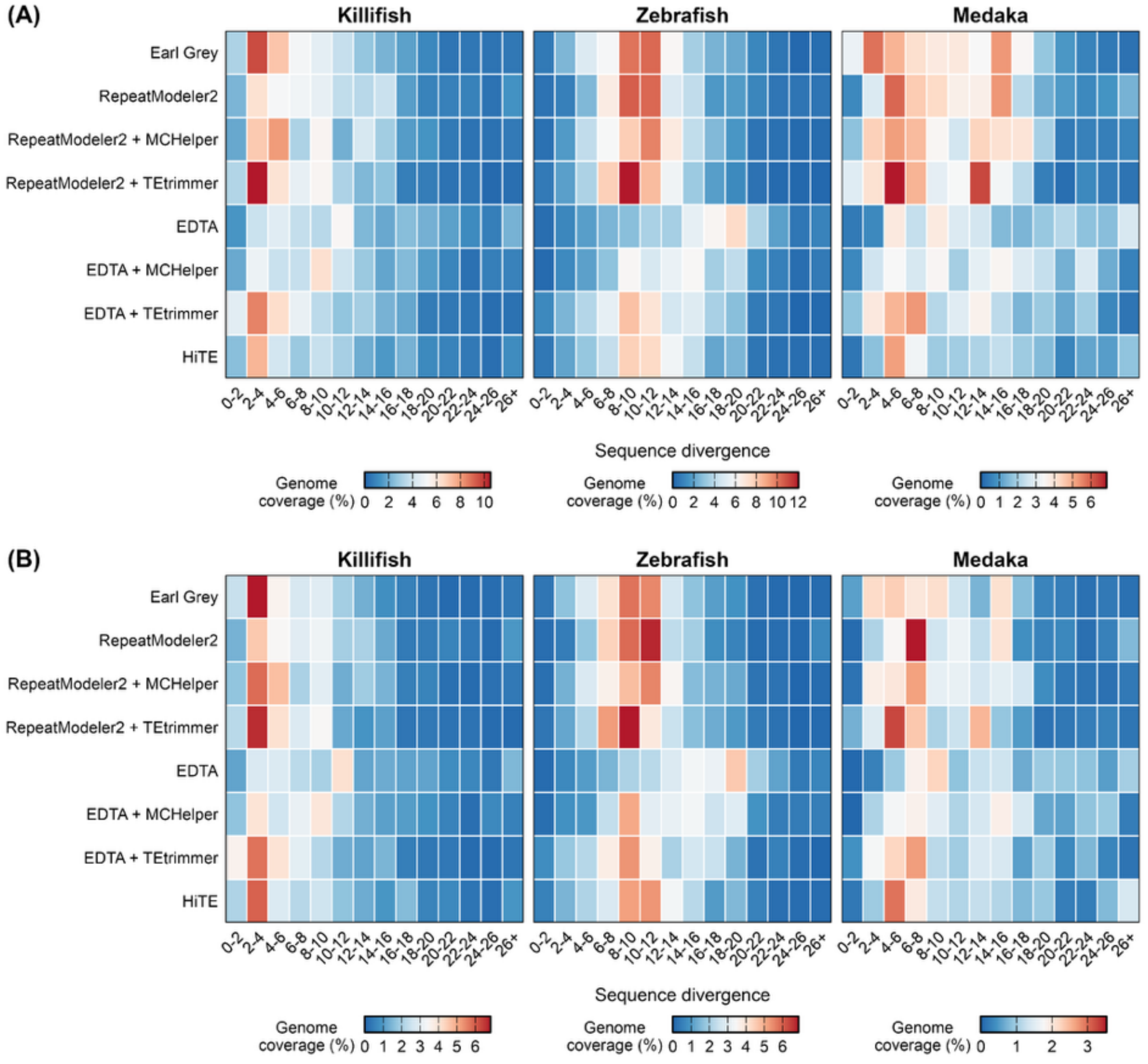


**Figure S5. Comparison of TE sequence divergence profiles across consensus libraries.**

For each species, rows represent TE consensus libraries and columns represent sequence divergence bins. Divergence values were summarized as the length-weighted mean for each TE consensus sequence. Divergence bins cover 0–2%, 2–4%, …, and 24–26%, with values of 26% or higher grouped into a “26+” bin. Genome coverage of TE-derived regions was calculated for each divergence bin. Panel (A) shows all TE-derived regions identified using each library. Panel (B) shows only those with at least 50% of their length overlapping merged intervals defined across all eight libraries (Methods).

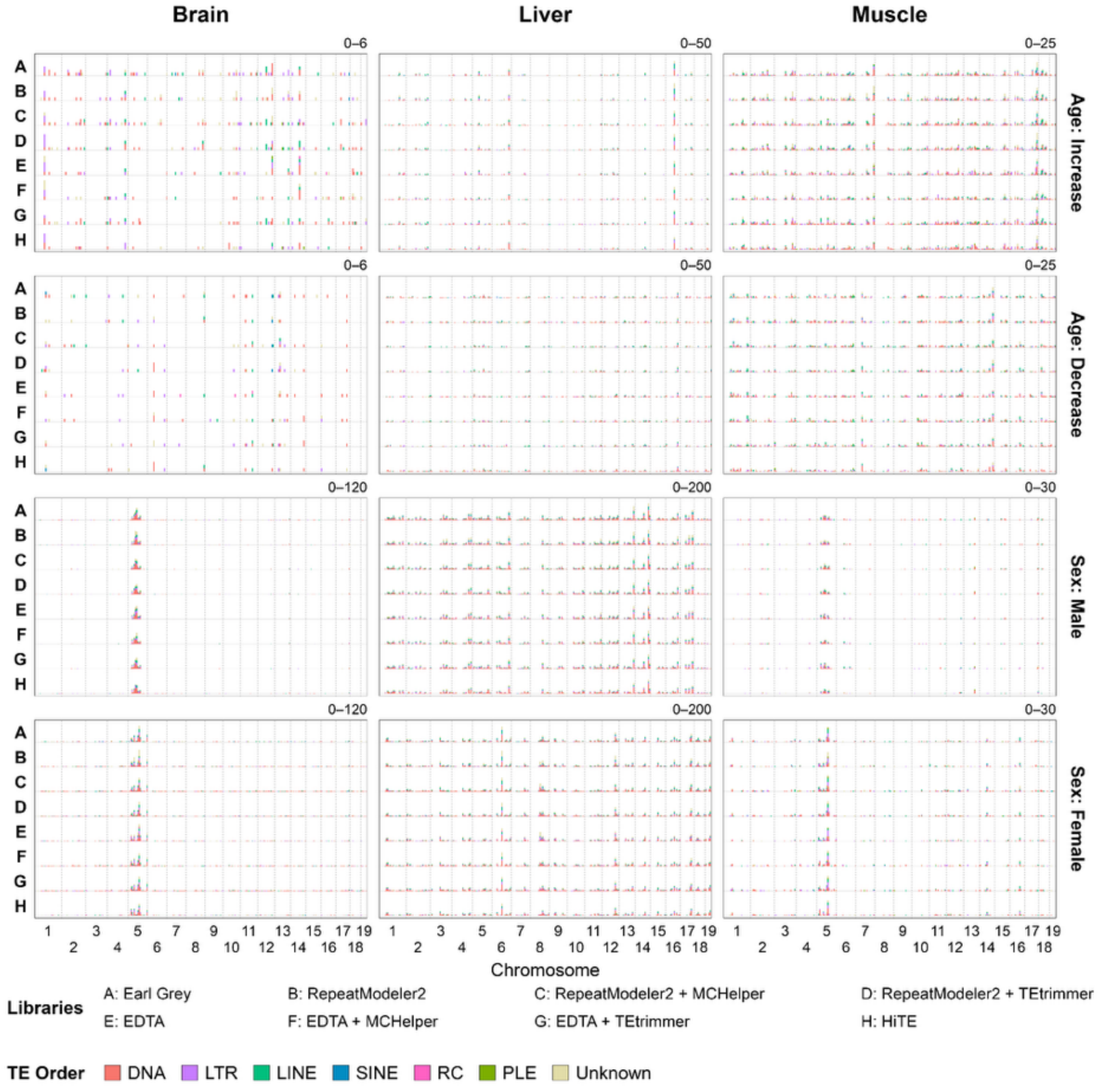


**Figure S6. Chromosomal distribution of age-associated and sex-associated TE loci across TE libraries.**

Locus selection criteria, plot format, and color coding are as in Figure 7D and E. Count scales are shared across libraries within each tissue and analysis type.

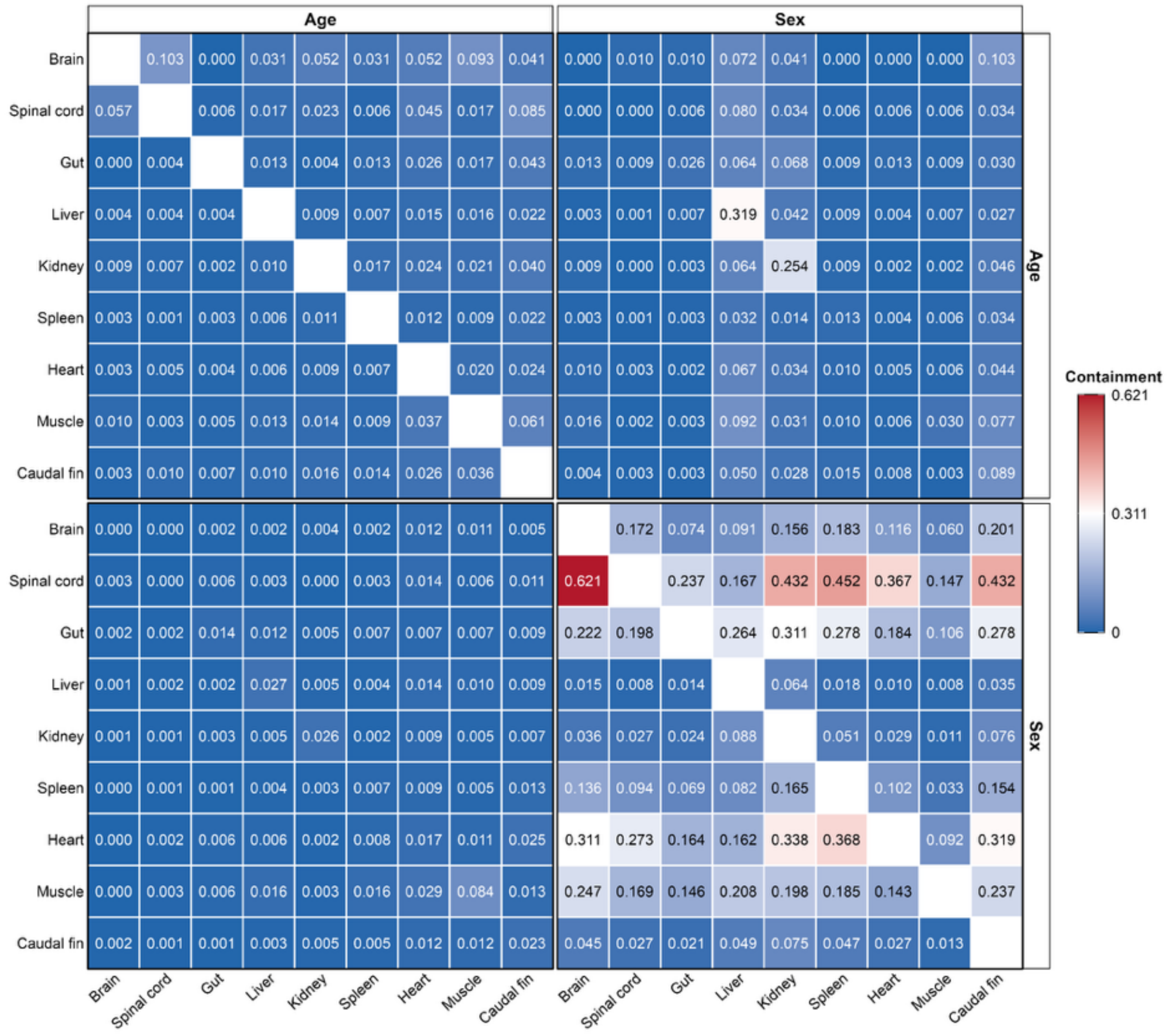


**Figure S7. Cross-tissue overlap of age- and sex-associated TE locus sets.**

Age- and sex-associated TE loci were identified using TE intervals defined by the Earl Grey–ULTRA workflow. Each cell shows the fraction of loci in the row set that are also present in the column set. This fraction was calculated as the number of shared loci divided by the total number of loci in the row set. Values are shown to three decimal places.

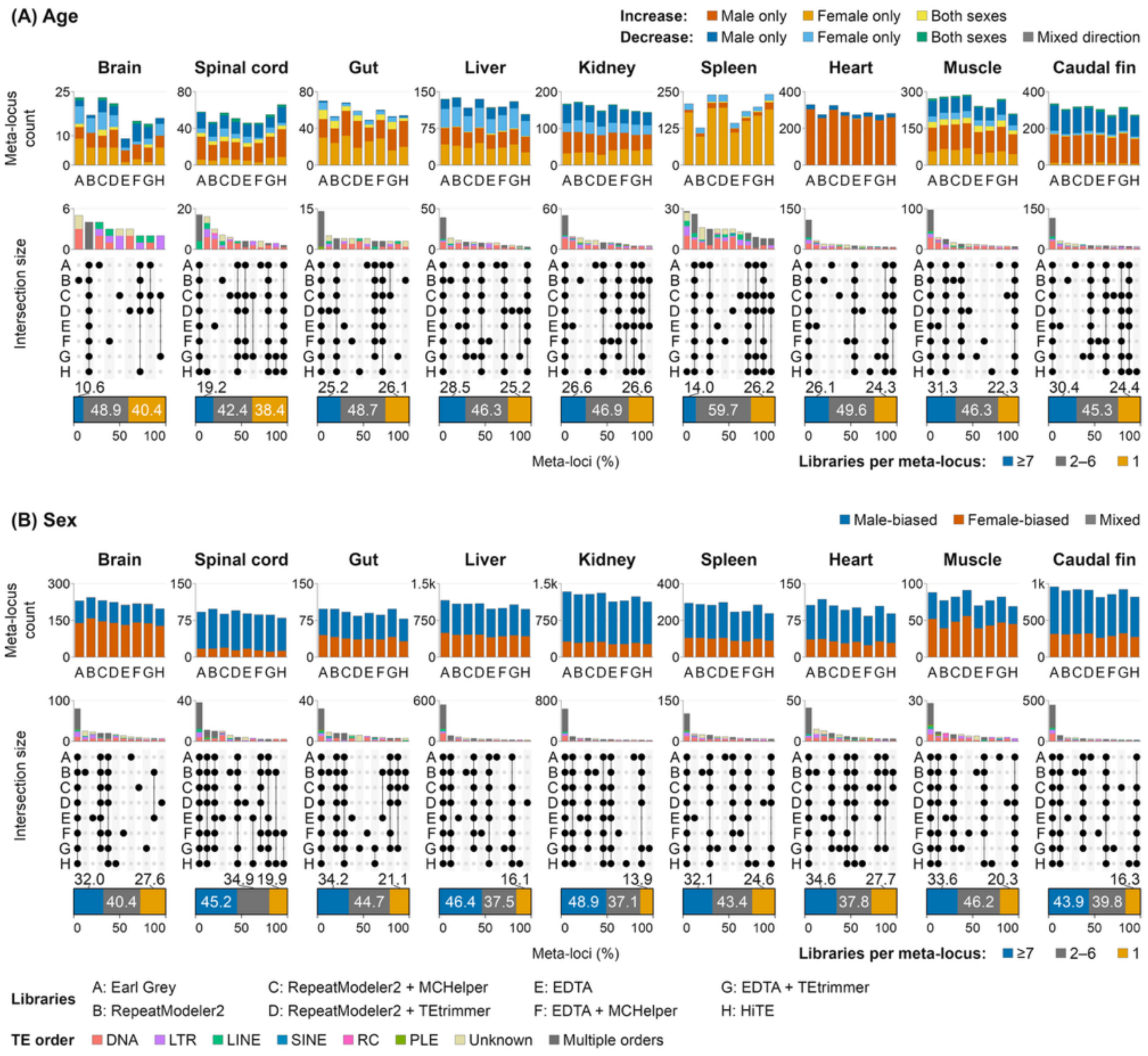


(C) Age-associated TE meta-loci

| | Brain | Spinal cord | Gut | Liver | Kidney | Spleen | Heart | Muscle | Caudal fin |
|---|---|---|---|---|---|---|---|---|---|
| Earl Grey | 1 | 2 | 1 | 2 | 2 | 4 | 1 | 4 | 1 |
| RepeatModeler2 | 4 | 6 | 7 | 1 | 1 | 8 | 6 | 3 | 6 |
| RepeatModeler2 + MCHelper | 2 | 3 | 2 | 7 | 4 | 2 | 2 | 2 | 3 |
| RepeatModeler2 + TEtrimmer | 3 | 5 | 4 | 3 | 6 | 3 | 3 | 1 | 2 |
| EDTA | 8 | 7 | 8 | 6 | 3 | 7 | 7 | 6 | 5 |
| EDTA + MCHelper | 6 | 8 | 3 | 5 | 5 | 6 | 4 | 7 | 8 |
| EDTA + TEtrimmer | 7 | 4 | 6 | 4 | 7 | 5 | 8 | 5 | 4 |
| HiTE | 5 | 1 | 5 | 8 | 8 | 1 | 5 | 8 | 7 |

(D) Sex-associated TE meta-loci

| | Brain | Spinal cord | Gut | Liver | Kidney | Spleen | Heart | Muscle | Caudal fin |
|---|---|---|---|---|---|---|---|---|---|
| Earl Grey | 3 | 3 | 1 | 1 | 1 | 2 | 2 | 2 | 1 |
| RepeatModeler2 | 1 | 1 | 3 | 3 | 4 | 3 | 1 | 6 | 5 |
| RepeatModeler2 + MCHelper | 2 | 5 | 4 | 4 | 3 | 5 | 3 | 4 | 3 |
| RepeatModeler2 + TEtrimmer | 4 | 2 | 7 | 2 | 2 | 1 | 6 | 1 | 4 |
| EDTA | 7 | 4 | 5 | 7 | 7 | 7 | 5 | 7 | 8 |
| EDTA + MCHelper | 5 | 6 | 6 | 6 | 6 | 6 | 8 | 5 | 6 |
| EDTA + TEtrimmer | 6 | 7 | 2 | 5 | 5 | 4 | 4 | 3 | 2 |
| HiTE | 8 | 8 | 8 | 8 | 8 | 8 | 7 | 8 | 7 |

**Figure S8. Age- and sex-associated intergenic TE meta-loci identified using Telescope's locus-unique counts.**

Plot format and color coding are as in Figure 8.

**Table S1. Genome assemblies used in the present study**

| Species | Genome assembly accession | Assembly name/version |
|---|---|---|
| zebrafish | GCF_049306965.1 | GRCz12tu |
| medaka | GCA_053564925.1 | Olatipes_Hd-rR_3.1 |
| African turquoise killifish | GCF_043380555.1 | NfurGRZ-RIMD1 |
| African turquoise killifish[a] | GCA_014300015.1 | MPIA-NFZ-2.0 |
| African turquoise killifish[a] | GCF_001465895.1 | Nfu_20140520 |
| African turquoise killifish[a] | GCA_000878545.1 | NotFur1 |
| African turquoise killifish[a] | GCF_027789165.1 | UI_Nfuz_MZM_1.0 |
| human | GCF_009914755.1 | T2T-CHM13v2.0 |
| mouse | GCA_050437135.1 | T2T_mhaESC_v1.0 |

[a] Previously published African turquoise killifish assemblies were used for comparison with NfurGRZ-RIMD1.

**Data S1. Age- and sex-associated TE loci and their nearest genes in killifish.**
ZIP archive containing a CSV file of library-specific TE loci identified in the age-associated and sex-differential-expression analyses. Each row represents a library-specific genomic TE locus for a given analysis, tissue, and Telescope count mode. Column definitions are provided as a separate file.
