## Supplementary material for "Near-complete genome assembly of the African turquoise killifish enables locus-level profiling of transposable element expression": Data S1: Data_S1_column_definitions.txt

(1) analysis: Association analysis. age denotes the age-correlation analysis, and sex denotes the stage-adjusted male-versus-female differential-expression analysis.
(2) tissue: Tissue in which the association was identified: Brain, Spinal cord, Gut, Liver, Kidney, Spleen, Heart, Muscle, or Caudal fin.
(3) library: TE library workflow used to define the source locus: Earl Grey, RepeatModeler2, RepeatModeler2 + MCHelper, RepeatModeler2 + TEtrimmer, EDTA, EDTA + MCHelper, EDTA + TEtrimmer, or HiTE.
(4) mapping_mode: Telescope count mode. unique uses unique_count, representing reads assigned to a single annotated locus. multi uses TE_counts, including unique and ambiguous reads, with the reassignment mode set to average.
(5) te_id: Identifier of the original genomic TE source locus in the specified library. This identifier refers to an individual genomic locus rather than a consensus sequence, gene, or TE family and should be interpreted together with library.
(6) meta_locus_id: Identifier of a cross-library TE meta-locus.
(7) seqname: UCSC-style name of the reference sequence containing the TE locus, including chromosome and unplaced-scaffold names.
(8) refseq_sequence_id: Versioned RefSeq accession for the reference sequence specified by seqname, as defined in the reference chromosome-alias table.
(9) start_1based: Start position of the TE interval on the specified reference sequence, using 1-based inclusive coordinates.
(10) end_1based: End position of the TE interval, using 1-based inclusive coordinates. The interval length is end_1based − start_1based + 1 bp.
(11) strand: Strand of the source TE annotation, encoded as + or −. The nearest transcription start site (TSS) is identified independently of the TE strand.
(12) te_class: Repeat-class label retained from the source annotation: DNA (DNA transposons), LTR (long terminal repeat retrotransposons), LINE (long interspersed nuclear elements), SINE (short interspersed nuclear elements), RC (rolling-circle elements), PLE (Penelope-like elements), or Unknown (unresolved classification).
(13) te_family: TE family or classification string retained from the library-specific source annotation. Records represent individual genomic loci and are not aggregated by family.
(14) genomic_context: Genomic context, classified as Genic or Intergenic, inherited from the source TE-locus annotation. Both categories are included. This classification is separate from the nearest-TSS assignment.
(15) female_spearman_rho (age only): Spearman’s correlation coefficient between normalized TE expression and age in females. Positive and negative values indicate tendencies toward increasing and decreasing expression with age, respectively.
(16) female_spearman_p (age only): Two-sided P value for the female Spearman correlation, before multiple-testing correction.
(17) female_q_bh (age only): Benjamini–Hochberg-adjusted female Spearman P value.
(18) male_spearman_rho (age only): Spearman’s correlation coefficient between normalized TE expression and age in males. Positive and negative values indicate tendencies toward increasing and decreasing expression with age, respectively.
(19) male_spearman_p (age only): Two-sided P value for the male Spearman correlation, before multiple-testing correction.
(20) male_q_bh (age only): Benjamini–Hochberg-adjusted male Spearman P value.
(21) wald_pvalue_nominal (sex only): Two-sided DESeq2 Wald P value for the unshrunken male-versus-female coefficient.
(22) wald_padj_BH (sex only): Benjamini–Hochberg-adjusted wald_pvalue_nominal.
(23) log2FoldChange_M_vs_F (sex only): Unshrunken, stage-adjusted log2 fold change in TE expression in males relative to females. Positive and negative values indicate male- and female-biased expression, respectively.
(24) lfcSE (sex only): Standard error of the unshrunken log2FoldChange_M_vs_F estimate.
(25) shrunken_log2FoldChange_apeglm (sex only): apeglm-shrunken estimate of the stage-adjusted male-versus-female log2 fold change. Positive and negative values indicate male- and female-biased expression, respectively.
(26) nearest_tss_distance_bp: Minimum unsigned distance, in base pairs, between the TE interval and an annotated transcript TSS on the same reference sequence. For a TSS at coordinate t, the distance is calculated as max(start_1based − t, t − end_1based, 0). A value of 0 indicates that the TE interval overlaps a TSS. NA indicates that no eligible annotated TSS is present on the reference sequence.
(27) nearest_gene_id: NCBI GeneID at each equally nearest gene/TSS/strand site. 
(28) nearest_gene_symbol: RefSeq gene symbol at each nearest site, with original LOC-number symbols retained.
(29) nearest_gene_description: RefSeq gene description at each nearest site, including descriptions of LOC-number genes.
(30) nearest_gene_biotype: RefSeq gene_biotype at each nearest site, such as protein_coding or a noncoding-gene biotype.
(31) nearest_gene_strand: Strand (+ or −) of the annotated gene/transcript at each nearest site.
(32) nearest_transcript_id: RefSeq transcript accession(s) supporting each nearest TSS site.
Nearest-TSS fields: When multiple TSS sites are tied for the minimum distance, values for distinct sites are separated by pipes (|) and are listed in the same order across nearest-TSS-related columns. Multiple transcript accessions at the same TSS site are separated by semicolons (;). Literal pipe characters in annotation text are encoded as %7C.
